# Distinct execution modes of a biochemical necroptosis model explain cell type-specific responses and variability to cell-death cues

**DOI:** 10.1101/2022.02.25.481705

**Authors:** Geena V. Ildefonso, Marie Oliver-Metzig, Alexander Hoffmann, Leonard A. Harris, Carlos F. Lopez

## Abstract

Necroptosis is a form of regulated cell death that has been associated with degenerative disorders, autoimmune processes, inflammatory diseases, and cancer. To better understand the biochemical mechanisms of necroptosis cell death regulation, we constructed a detailed biochemical model of tumor necrosis factor (TNF)-induced necroptosis based on known molecular interactions. Intracellular protein levels, used as model inputs, were quantified using label-free mass spectrometry, and the model was calibrated using Bayesian parameter inference to experimental protein time course data from a well-established necroptosis-executing cell line. The calibrated model accurately reproduced the dynamics of phosphorylated mixed lineage kinase domain-like protein (pMLKL), an established necroptosis reporter. A dynamical systems analysis identified four distinct modes of necroptosis signal execution, which can be distinguished based on rate constant values and the roles of the deubiquitinating enzymes A20 and CYLD in the regulation of RIP1 ubiquitination. In one case, A20 and CYLD both contribute to RIP1 deubiquitination, in another RIP1 deubiquitination is driven exclusively by CYLD, and in two modes either A20 or CYLD acts as the driver with the other enzyme, counterintuitively, inhibiting necroptosis. We also performed sensitivity analyses of initial protein concentrations and rate constants and identified potential targets for modulating necroptosis sensitivity among the biochemical events involved in RIP1 ubiquitination regulation and the decision between complex II degradation and necrosome formation. We conclude by associating numerous contrasting and, in some cases, counterintuitive experimental results reported in the literature with one or more of the model-predicted modes of necroptosis execution. Overall, we demonstrate that a consensus pathway model of TNF-induced necroptosis can provide insights into unresolved controversies regarding the molecular mechanisms driving necroptosis execution for various cell types and experimental conditions.

## INTRODUCTION

Apoptosis is widely recognized as the primary form of programmed cell death, characterized by a concerted dismantling of the cell into apoptotic bodies that can be easily processed by the immune system.^1^ Conversely, necroptosis is an alternative form of programmed cell death in which the cell membrane is ruptured, leading to immune response activation.^2,3^ Various human diseases, including neurodegenerative disorders and cancer, have been associated with necroptosis.^4^ Induction of necroptosis is also currently being explored as an alternative anticancer therapy, since apoptosis resistance is a hallmark of cancer.^5–7^ Although many of the primary molecular species involved in necroptosis have been identified,^8^ including receptor interacting protein kinase-1 (RIP1), RIP3, and mixed lineage kinase domain-like protein (MLKL), efforts to target necroptosis dysregulation or leverage it therapeutically are hindered by the lack of a detailed, mechanistic understanding of the biochemical pathways driving necroptosis execution.^4^

Prior studies^9–16^ of necroptosis identified multiple mechanisms of ubiquitination regulation, including K63, K48, and M1 chains, which lead to phosphorylation of RIP1 and RIP3, phosphorylation and activation of cell death marker MLKL,^9^ and plasma membrane permeabilization resulting in cell death.^8^ The K63-specific deubiquitinase CYLD^17^ (cylindromatosis lysine 63 deubiquitinase) and the ubiquitin-editing enzyme A20^15^ (tumor necrosis factor, alpha-induced protein 3) are both known to mediate deubiquitination of RIP1, which precedes RIP1 phosphorylation, by cleaving K63 ubiquitin chains and facilitating the formation of complex II.^10–16^ Therefore, both enzymes are generally considered drivers of necroptosis.^18^ However, CYLD- and A20-driven deubiquitination of RIP1 have been variously reported as pro- and anti-necroptotic in different cell types: some studies have shown that CYLD drives RIP1 deubiquitination,^12,17,19,20^ while others have implicated A20^21–23^ or reported equal contributions from both enzymes.^24–26^ These varying reports have led to unresolved controversies within the field regarding the specific molecular mechanisms of complex II formation and subsequent necroptotic cell death.^4^ For example, Vanlangenakker et al.^26^ showed that repression of CYLD in L929 cells, a murine fibrosarcoma cell line, protects from tumor necrosis factor (TNF)-induced necroptosis but, unexpectedly, A20 repression increases sensitivity to necroptosis. A recent time-resolved analysis of necroptosis rates and network components revealed an incoherent feedforward loop through which NF-κB and A20 counteract pro-necroptotic signaling in L929 cells,^27^ but it remains unclear how general or cell context-dependent this regulatory control of necroptosis is.

Here, we present, to our knowledge, the first detailed biochemical model of TNF-induced necroptosis. The model is derived from published literature and incorporates known biology obtained from decades’ worth of experimental studies (Table 1). We calibrate the model to experimental phosphorylated MLKL (pMLKL) time course data from TNF-treated mouse fibrosarcoma cells at multiple TNF doses. We then perform a dynamical systems analysis that identifies four modes of necroptosis signal execution. In one case, A20 and CYLD contribute approximately equally to RIP1 deubiquitination, such that both must be knocked out to delay necroptosis induction (knocking out one has no effect, since the signal can be rerouted through the other). In another, RIP1 deubiquitination is driven exclusively by CYLD, with A20 being effectively inactive. In the other two modes, either A20 or CYLD acts as the driver of RIP1 deubiquitination, with the other enzyme, counterintuitively, acting to inhibit necroptosis (consistent with the observation by Vanlangenakker et al.^26^). We also perform sensitivity analyses to identify proteins and kinetic parameters that can be targeted within each mode to modulate pMLKL dynamics and time-to-death (TTD) by necroptosis. We find that, for two modes, proteins and rate constants centered around RIP1 ubiquitination regulation in complex I have the most significant effect on necroptosis execution. For the other two, potential targets include factors involved in the balance between complex II degradation and necrosome formation. Overall, our results show that a consensus pathway model of TNF-induced necroptosis can explain numerous experimentally observed behaviors, including conflicting and counterintuitive results from multiple studies involving different cell types. Following a detailed description of our proposed model, we present results of the parameter calibration, dynamical systems analysis, *in silico* knockout experiments, and sensitivity analyses. We conclude with a discussion of the broader implications of our results, including important insights into the molecular mechanisms of necroptosis execution and the potential for using the model to identify novel pro- and anti-necroptosis therapeutic targets.

**Table 1:** Key proteins involved in necroptosis.

| Protein | Role in necroptosis | References |
| --- | --- | --- |
| A20 | Ubiquitin-editing enzyme responsible for deubiquitinating RIP1 in complex I | 21,22 |
| Caspase-8 | Heterodimerizes with cFLIPL (long isoform), leading to cleavage and inactivation of RIP1 and RIP3 in complex II | 38,97 |
| cFLIPL | Heterodimerizes with caspase-8, leading to cleavage and inactivation of RIP1 and RIP3 in complex II | 39,98 |
| cIAP1/2 | Catalyzes, via its RING domains, the activating K63-linked polyubiquitination of RIP1 | 26,69 |
| CYLD | Deubiquitinates RIP1 in either complex I or within the RIP1-RIP3 necrosome | 12,17 |
| FADD | TNFR1-interacting scaffold protein in complex II | 18,99,100 |
| LUBAC | TNFR1-interacting protein recruited by cIAP1/2 in complex I that promotes RIP1 ubiquitination | 26,70 |
| MLKL | Recruited to the necrosome by RIP1, where it is phosphorylated, leading to cell death by membrane rupture | 44,101,102 |
| RIP1 | A multifunctional adaptor protein in the necrosome that recruits and activates RIP3 and MLKL | 26,103,104 |
| RIP3 | Recruited to the necrosome by binding to and cross-phosphorylating RIP1 | 101,105,106 |
| TNF | Pleiotropic pro-inflammatory cytokine that activates necroptosis in the absence of caspase activity | 107 |
| TNFR1 | TNF receptor superfamily member death receptor that recruits RIP1 to complex 1 | 30,108 |
| TRADD | TNFR1-interacting protein in complexes I and II that serves as a docking adaptor for the binding of RIP1 to TRAF2 | 32,108 |
| TRAF2 | TNFR1-interacting protein that recruits cIAP1/2 to complex I, promoting K63-linked RIP1 ubiquitination | 109,110 |

## RESULTS

### A biochemical model of TNF-induced necroptosis describes the formation of key signaling complexes along the path to cell death

The death receptor ligand TNF,^28^ an extensively studied inducer of necroptosis and well-known master regulator of inflammation, has been at the forefront of numerous fundamental discoveries concerning the interplay between cell death and survival pathways.^26^ Here, we propose a detailed, mechanistic model of TNF-induced necroptosis based on an extensive review of the literature (Table 1, with references). The model comprises 14 proteins interacting via 40 reactions (all mass action) to produce 37 biochemical species, including complex I, complex II, and the necrosome (Fig. 1), three key macromolecular complexes along the path from cell-death cue to necroptosis execution. Below, we describe in detail the steps involved in the formation of each complex, beginning with TNF binding to TNF receptor 1 (TNFR1) and ending at phosphorylation of the necroptosis cell death reporter MLKL. A model schematic is provided as a visual aid (Fig. 1), with reactions, including association, dissociation, phosphorylation, ubiquitination, deubiquitination, and degradation, denoted as “*R_N_*,” where *N* is the reaction number. Note that protein synthesis is omitted from the model because all experiments were performed in the presence of cycloheximide (see Materials and Methods), commonly used to sensitize cells to TNF.^29^

**Figure 1:**
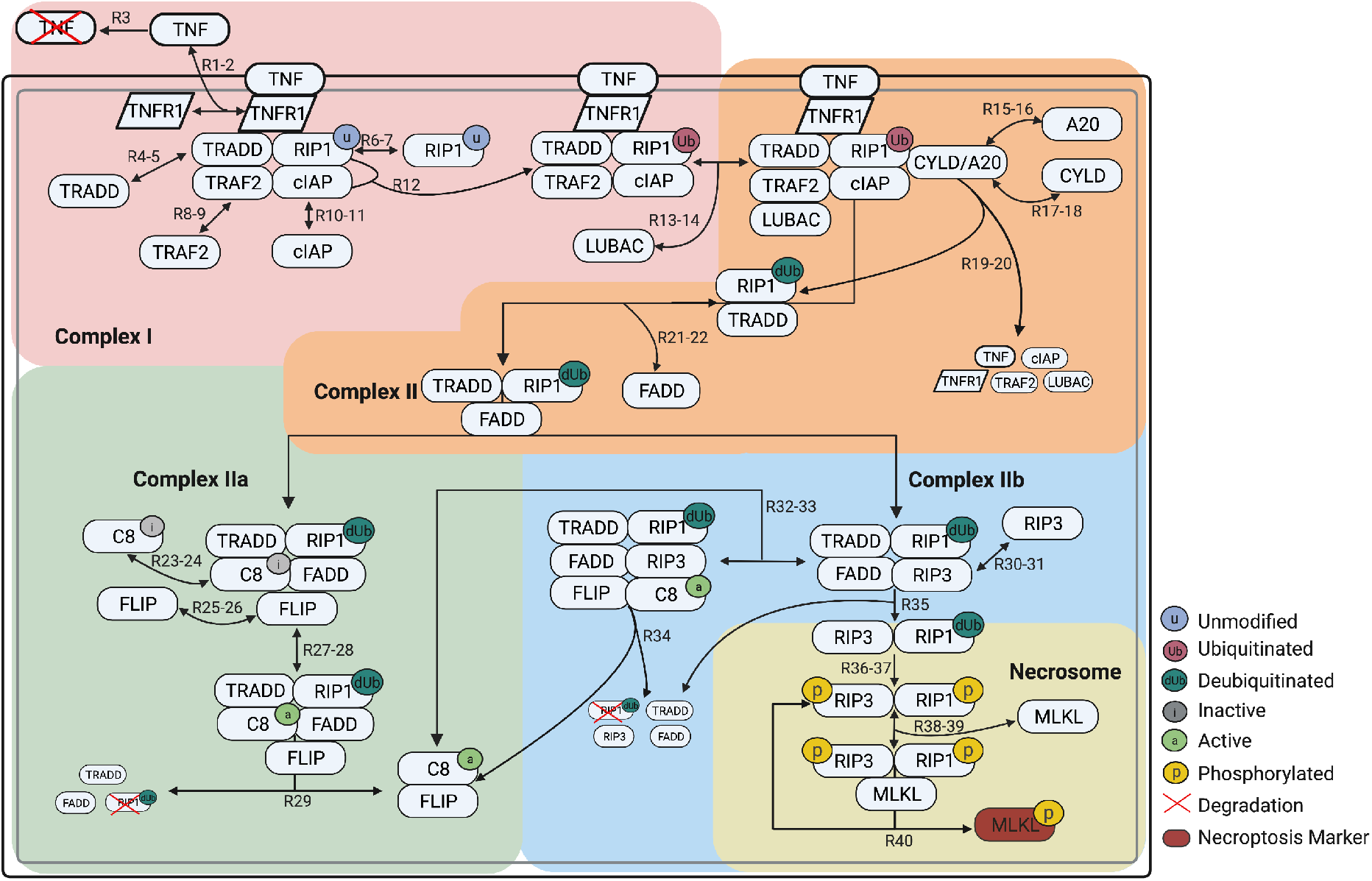
Schematic of the necroptosis execution model. The diagram is color coded to highlight the processes involved in formation of complex I, complex II, complex Ila, complex Ilb, and the necrosome. Arrows are labeled with ‘*R_N_*’ or ‘*R_N-M_*, where *N* and *M* correspond to reaction indices in the model. In many cases (but not all; see text), ‘*R_N-M_*’ denotes a set of reversible reactions, with *N* the index of the forward direction and *M* the index of the reverse. Note that unmodified (u) and deubiquitinated (dUb) RIP1 are considered distinct states and are involved in different reactions. Created with BioRender.com.

Signaling through the necroptosis pathway is initiated when the cytokine TNF binds to the extracellular domain of TNFR1 (*R*_1-2_), which protects TNF from degradation (*R*_3_) and activates the receptor by causing a conformational change in its intracellular domain.^26,30,31^ The adaptor protein TRADD (TNFR1-associated death domain) is then recruited to the intracellular domain of TNFR1 (*R*_4-5_) to facilitate binding of RIP1 (unmodified; *R*_6-7_) and TRAF2 (TNFR-associated factor 2; *R*_8-9_).^32–34^ TRAF2 recruits and binds cIAP1/2 (cellular inhibitor of apoptosis proteins 1 and 2; *R*_10-11_), which add non-degradative polyubiquitin chains to RIP1 (*R*_12_).^9^ Ubiquitinated RIP1 recruits other necessary components to the complex, including LUBAC (linear ubiquitin chain assembly complex; *R*_13-14_). We refer to the supramolecular structure, which is anchored to the cell membrane and composed of TNF, TNFR1, TRADD, ubiquitinated RIP1, TRAF2, cIAP1/2, and LUBAC, as complex I^29,35^ (Fig. 1, *pink*). Biologically, complex I is known to drive multiple pathways in addition to necroptosis, including apoptosis and the inflammatory NF-κB pathway.^36^

Formation of complex I is followed by deubiquitination of RIP1 by the enzymes A20^15,21^ and CYLD,^12,17,19,20^ which competitively bind to RIP1 in its ubiquitinated state (*R*_15-18_), causing cleavage, deubiquitination, and release in association with TRADD and the dissolution of complex I (*R*_19-20_). The RIP1:TRADD heterodimer then recruits FADD (Fas-associated protein with death domain; *R*_21-22_), initiating the formation of complex II, also known as the cytosolic death-inducing signaling complex (Fig. 1, *orange*). Complex II can then be modified via two competing paths, one anti-necroptotic and one pro-necroptotic. The anti-necroptotic path involves FADD, via its death effector domain, mediating the recruitment of inactive Caspase 8 (C8i; *R*_23-24_),^37^ which subsequently binds FLIP (cellular FADD-like IL-1β-converting enzyme-inhibitory protein; *R*_25-26_), resulting in the complex commonly referred to as complex IIa (Fig. 1, *green*).^26,37^ FLIP then oligomerizes with C8i to produce active Caspase-8 (C8a; *R*_27-28_),^38,39^ which cleaves RIP1 for truncation (i.e., degradation), resulting in dissolution of the complex and release of the active C8a:FLIP heterodimer^40,41^ (*R*_29_) that directly inhibits necroptosis (*R*_32-34_; see below).

The pro-necroptotic path involves formation of complex IIb (Fig. 1, *blue*), which occurs when deubiquitinated RIP1 in complex II recruits RIP3 (receptor-interacting protein kinase 3; *R*_30-31_), blocking C8i recruitment (*R*_23-24_). The C8a:FLIP heterodimer can then be recruited to complex IIb (*R*_32-33_), which cleaves RIP1 for truncation, leading to dissolution of the complex (*R*_34_). Alternatively, RIP3 and deubiquitinated RIP1 can dissociate from complex IIb as a heterodimer (*R*_35_).^26^ Cross-phosphorylation of RIP3 (*R*_36_) and then RIP1 (*R*_37_), followed by recruitment of MLKL (*R*_38-39_),^42,43^ results in the necroptosis signaling complex, known as the necrosome (Fig. 1, *yellow*).^26^ Phosphorylation of MLKL^44^ in the necrosome by phosphorylated RIP1 and RIP3 is followed by release of pMLKL from the phosphorylated RIP1:RIP3 heterodimer (*R*_40_), which is again free to bind MLKL. We assume dephosphorylation and degradation of the phosphorylated RIP1:RIP3 heterodimer is negligible, consistent with experimental reports.^45^ Translocation of pMLKL to the cell membrane^46^ then causes rapid plasma membrane rupture and inflammatory response due to the release of DAMPs (damage-associated molecular patterns) and cytokines,^47^ ultimately resulting in cell death.

### Western blots and mass spectrometry enable Bayesian parameter estimation of the necroptosis model

To explore the dynamics of our computational necroptosis model, we first calibrated it to experimental protein time course data using a Bayesian parameter estimation approach.^48^ Briefly, we used L929 cells, a murine fibrosarcoma cell line that is a well-established model system for studying necroptosis.^26^ Cells were treated with 100, 10, 1, and 0.1 ng/ml of TNF over 16 hours and pMLKL levels were estimated at multiple time points via Western blot using densitometry (Fig. 2A). To quantify initial protein abundances, used as inputs to the model, we used label-free mass spectrometry in untreated L929 cells for proteins C8, FADD, MLKL, RIP3, TRADD, and TRAF2 (Fig. 2B). All other initial protein levels (other than TNF, which depends on applied dose) were set to values based on biologically plausible assumptions (Supplementary Table S1). Parameter estimation was then performed using PyDREAM^48^ (Fig. 2C), a multi-chain Monte Carlo sampling tool, with a multi-objective cost function that included data from the two highest TNF doses (100 and 10 ng/ml; Supplementary Fig. S1). In all, an ensemble of 10,628 parameter sets was obtained (Supplementary Fig. S2), all of which reproduce the experimental data reasonably well^49^ (see Materials and Methods for additional details). Model simulations at the two lowest TNF doses (1 and 0.1 ng/ml; Fig. 2C) showed good correspondence to experimental data, providing a simple validation of the model fits.

**Figure 2:**
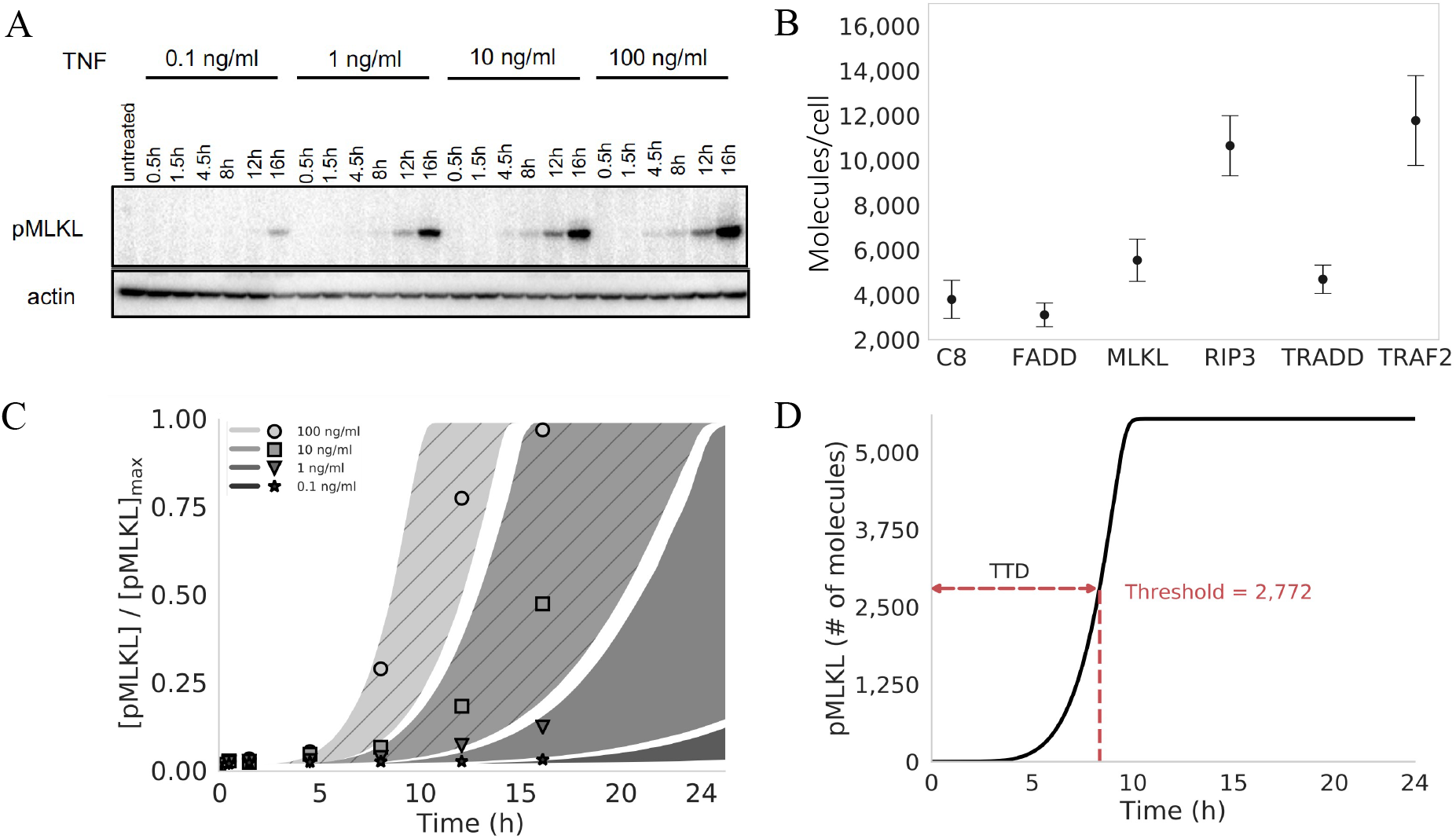
Proteomics, parameter calibration, and time-to-death. (A) Western blots for phosphorylated MLKL (pMLKL) at multiple time points in L929 (murine fibrosarcoma) cells under 0.1–100 ng/ml TNF stimulation. Actin, used as a loading control, is also shown for comparison. (B) Mass spectrometry data from untreated L929 cells for multiple proteins involved in necroptosis execution. Points represent the median of three replicates (used as input to the computational model); error bars span the interquartile range. (C) Simulated pMLKL time courses (plotted as 95% probability envelopes) for 0.1–100 ng/ml TNF stimulation (same concentrations as in *A)* based on 10,628 parameter sets obtained using Bayesian parameter estimation. The model was calibrated to the 100 and 10 ng/ml TNF data only (shaded regions with diagonal lines); time courses for the lowest TNF concentrations (shaded regions with no diagonal lines) amount to a simple model validation. Points correspond to the Western blot data in *A*, quantified via densitometry. Points and shaded regions are colored the same, based on TNF dose. (D) Illustration of the time-to-death (TTD) metric used to quantify cell death *in silico.* A hard threshold of 2,772 molecules (half the median MLKL level in *B)* was chosen to signify cell death (see Materials and Methods). MLKL: mixed lineage kinase domain-like protein; TNF: tumor necrosis factor.

### A dynamical systems analysis identifies four distinct necroptosis execution modes differing by mechanism of RIP1 ubiquitination regulation

We performed a dynamical systems analysis to explore the possibility that distinct “modes of necroptosis execution” exist within the parameter set ensemble obtained from Bayesian parameter estimation. The rationale is that while different parameterizations of the model achieve cell death at approximately equal times, they may arrive there via significantly different sequences of molecular events. We utilized a computational tool^50^ that identifies subnetworks of reactions that dominate the production or consumption of a target species, pMLKL in this case, at user-specified times along a time course. Each subnetwork is given an integer label and each time point is associated with a subnetwork. Thus, a continuous concentration time course is “digitized” into a sequence of integers, which we refer to as a “dynamical signature.” This transformation enables simple comparisons between time courses obtained with different parameter sets using standard dissimilarity metrics, such as the longest common subsequence.^51^ Applying this approach to all 10,628 parameter sets obtained from Bayesian parameter estimation of our necroptosis model and clustering the resulting dynamical signatures using a spectral clustering algorithm,^52^ we obtained four distinct clusters, or modes of necroptosis execution (Fig. 3A and Supplementary Fig. S3; see Materials and Methods for additional details).

**Figure 3:**
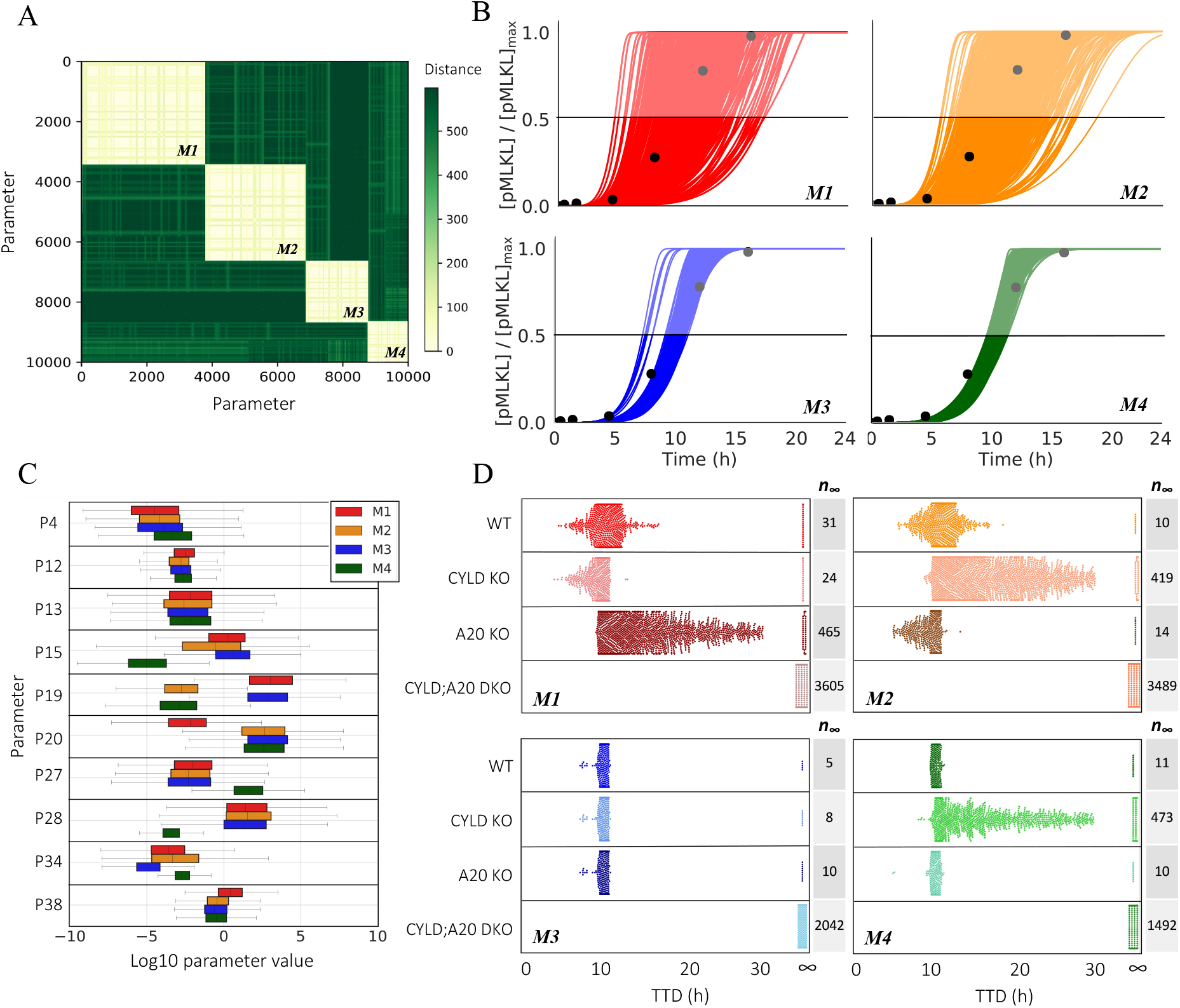
Four modes of necroptosis execution exhibit variability in temporal dynamics and differ in rate constant values and responses to CYLD and A20 knockouts. (A) Clustering analysis of simulated time courses (100 ng/ml TNF) from 10,628 parameter sets reveals four distinct modes of execution (*M1,* …, *M4*). Dissimilarity (“distance”) between dynamical signatures (digitized time courses) was quantified using the longest common subsequence (see Materials and Methods). (B) Simulated time courses (100 ng/ml TNF) of the necroptosis marker, phosphorylated MLKL (pMLKL), show significantly more variability in time-to-death (TTD; defined as the time at which pMLKL reaches its half-maximal value) in modes 1 and 2. Time courses for all parameter sets associated with each mode are shown. Experimental Western blot data (black circles; quantified from Fig. 2A) are included to illustrate the model fit for each mode. (C) Variations in the values of 10 rate constants (P*N*, where *N* corresponds to the associated reaction index in Fig. 1) distinguish the four modes of execution. Note that P12 and P13 are included because of the high sensitivity of the model to variations in their values (discussed in the next subsection). (D) Knockouts of CYLD and A20 (100 ng/ml TNF) differentially affect TTD, relative to wild type (WT), across the four modes of execution (each dot corresponds to a parameter set). Note that CYLD;A20 double knockout inhibits cell death in all cases (TTD = ∞). The number of parameter sets that do not result in cell death (*n*_∞_) are included for all modes under all conditions. KO: knockout; DKO: double knockout.

Interestingly, two of the execution modes exhibit significantly more variability in pMLKL temporal dynamics and TTD (defined in Fig. 2D) across their associated parameter sets than the other two (Fig. 3B). This suggests the modes harbor fundamental differences in rate constant values that lead to differential robustness to parameter variations. To explore this further, we compared the distributions of rate constants across modes and identified eight (out of 40) with significant differences (>7.5-fold) between the largest and smallest mean (Fig. 3C; additional distributions are shown in Supplementary Fig. S5). We also consider distributions for two rate constants (P12 and P13; parameter indices correspond to reaction indices in Fig. 1) with much smaller differences across means (~3-fold in both cases) but for which the model exhibits high sensitivity (discussed in the next subsection). In all, these 10 rate constants correspond to reactions spanning the model topology, starting with the association of TRADD to complex I (P4), which has a somewhat increased rate in mode 4. Further downstream, the rate constant for ubiquitination of RIP1 by cIAP (P12) is slightly larger in mode 1 than in the other modes. Small differences are also seen for the binding rate of LUBAC to complex I (P13). The rate constant for binding of A20 to ubiquitinated RIP1 (P15) is significantly smaller in mode 4 than in the other modes and somewhat smaller in mode 2 relative to modes 1 and 3. Deubiquitination of RIP1 by A20 (P19) is significantly reduced in modes 2 and 4, while, interestingly, the rate constant for RIP1 deubiquitination by CYLD (P20) in mode 1 is reduced by almost the same amount relative to the other modes. For activation/deactivation of C8 in complex IIa, which is a critical step in the pathway for determining whether the cell will progress to necroptosis, mode 4 has both a significantly larger activation (P27) and significantly smaller deactivation (P28) rate constant. The rate constant for subsequent RIP1 degradation by the active C8a:FLIP heterodimer to complex IIb (P34), which inhibits necroptosis, is somewhat smaller in mode 3 and larger in mode 4 relative to the other modes. Finally, the binding rate constant for MLKL to the phosphorylated RIP1:RIP3 heterodimer (P38), the final step in the formation of the necrosome, is somewhat increased in mode 1. These results clearly illustrate that significant differences exist in the values of rate constants across the modes of execution, despite the similarities in pMLKL temporal dynamics.

CYLD and A20 are known regulators of RIP1 deubiquitination^10–16^ but have been reported as both drivers and inhibitors of necroptosis in different cell types.^12,17,19,20,24–26^ To investigate the roles of CYLD and A20 in our necroptosis model, we performed *in silico* CYLD and A20 knockout (KO) experiments and compared TTD distributions to the unperturbed, i.e., “wild-type” (WT), case (Fig. 3D). Unsurprisingly, in all cases CYLD/A20 double KO (DKO) prevents cell death (TTD = ∞). However, for single CYLD KO and A20 KO, we see highly variable responses across the four modes of execution. For mode 1, we see that knocking out A20 leads to a general increase in TTD (i.e., decrease in necroptosis sensitivity) across the parameter sets, consistent with A20 acting as a regulator of RIP1 ubiquitination and driver of necroptosis.^15,21^ Conversely, CYLD KO results in a general reduction in TTD (i.e., increase in sensitivity), indicating that CYLD in mode 1 counterintuitively operates as an inhibitor of necroptosis. We see the opposite trends in mode 2: A20 KO reduces TTD, while CYLD KO leads to a general increase in TTD across the parameter sets. This result is consistent with observations by Vanlangenakker et al.^26^ that A20 depletion can sensitize cells to death by necroptosis. In mode 3, we see that single KOs of A20 and CYLD have no effect on TTD. Since DKO prevents cell death in all cases, this reveals that A20 and CYLD both drive RIP1 deubiquitination and, hence, when one enzyme is knocked out signal flow diverts through the other. Finally, in mode 4, CYLD KO leads to a general increase in TTD, like mode 2; however, A20 KO has no effect, as in mode 3. In all, the results of *in silico* KO experiments reveal distinct differences in the roles of A20 and CYLD in RIP1 ubiquitination regulation among the four model-predicted modes of necroptosis execution (summarized in Table 2).

**Table 2:** Roles of A20 and CYLD in RIP1 deubiquitination and necroptosis execution in the four signal execution modes. ⇓: decrease; ⇑: increase; ⇔: no change; TTD: time-to-death.

|  |  |  |
| --- | --- | --- |
| **Mode 1** | • A20 ↓ TTD ↑<br>• CYLD ↓ TTD ↓ | • A20 deubiquitinates RIP1<br>• CYLD (counterintuitively) inhibits necroptosis |
| **Mode 2** | • CYLD ↓ TTD ↑<br>• A20 ↓ TTD ↓ | • CYLD deubiquitinates RIP1<br>• A20 (counterintuitively) inhibits necroptosis |
| **Mode 3** | • CYLD ↓ TTD ↑<br>• A20 ↓ TTD ⇔ | • CYLD deubiquitinates RIP1<br>• A20 has no significant role in necroptosis execution |
| **Mode 4** | • A20 ↓ TTD ⇔<br>• CYLD ↓ TTD ⇔<br>• A20 ↓ CYLD ↓ TTD ↑ | • Both A20 and CYLD can drive RIP1 deubiquitination<br>• If one is knocked out, the signal can reroute through the other<br>• Double KO prevents cell death (true for all modes) |

### Ubiquitination of RIP1 by cIAP in complex I and binding of LUBAC to complex I are global modulators of necroptosis sensitivity across execution modes

Targeting necroptosis by small molecule modulators has emerged as a promising approach for both cancer therapy and treatment of inflammatory diseases.^53^ It is of interest, therefore, to determine if modulating factors exist that are common across all modes of execution, which could represent novel therapeutic targets. Towards this end, we performed sensitivity analyses based on “representative” parameter sets for each mode (automatically generated by our dynamical systems analysis tool;^50^ see Materials and Methods for details) over the 14 non-zero initial protein concentrations (Fig. 4A) and 40 rate constants (Fig. 5A, Supplementary Fig. S4). Initial protein concentrations were varied ± 20% around a reference set of concentrations (Supplementary Table S1) used for parameter estimation; rate constant values were varied ± 20% around the representative parameter set for each mode. We then validated the results of these analyses (i.e., to confirm they are not specific to the representative parameter set) by performing, for all parameter sets associated with each mode, *in silico* knockdowns (KDs) by 70% and 10-fold overexpressions (OEs) for the initial concentrations^54,55^ (Fig. 4B) and by varying the rate constants values ± 10-fold (Fig. 5B).

**Figure 4:**
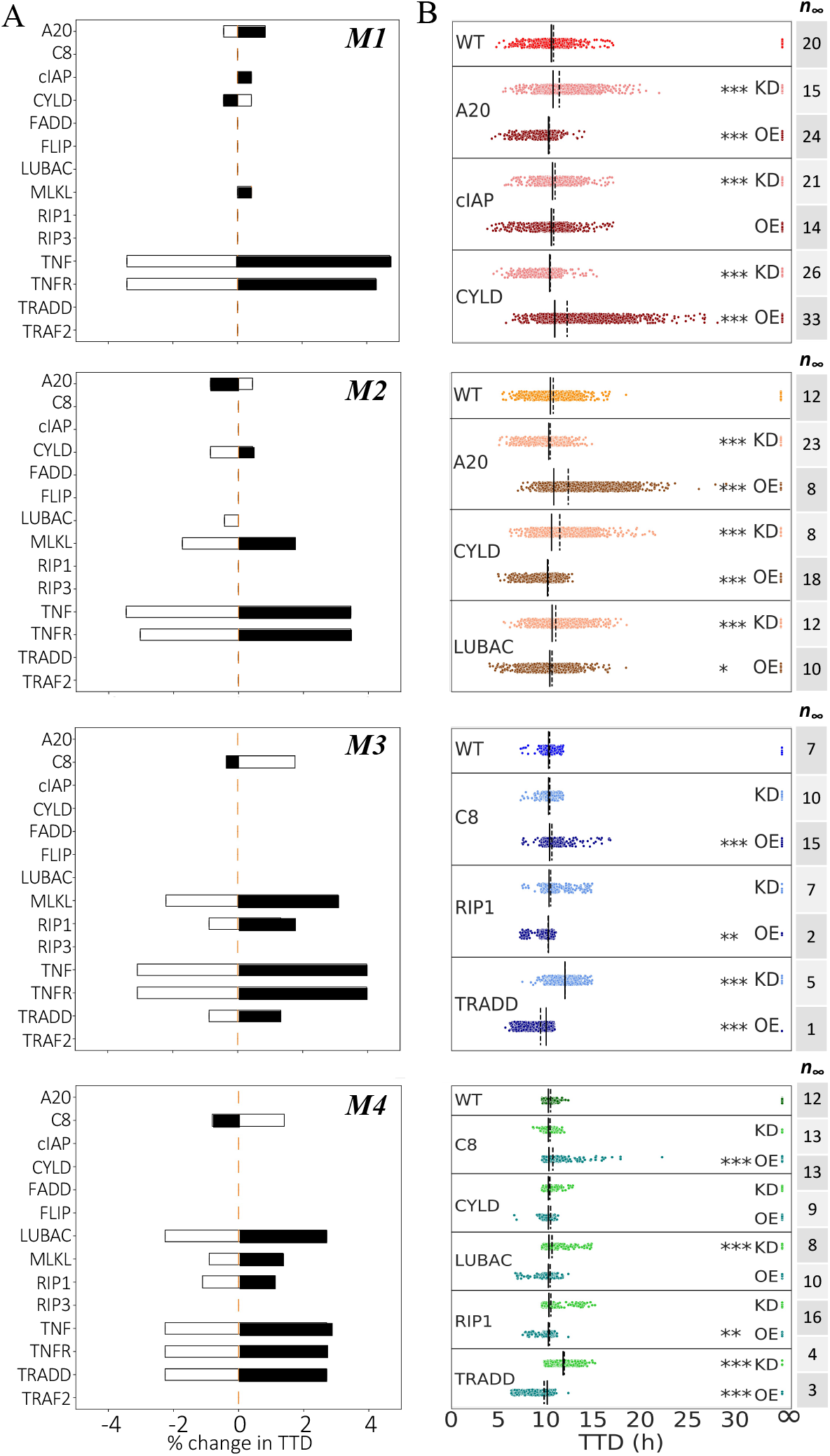
Sensitivity analyses and model-predicted protein targets for each mode of execution. (A) Changes in TTD for “representative” parameter sets of each mode. Black shaded regions signify decreases in initial protein concentrations; white shaded regions signify increases. (B) Knockdown (KD; 70%) and overexpression (OE; 10-fold) of potential targets identified in *A* for all parameter sets for each mode. The number of parameter sets that do not result in cell death (*n*_∞_) are included. Solid black lines = medians, dashed black lines = means; * p < 0.05, ** p < 0.01, *** p < 0.001 (Mood’s median test).

**Figure 5:**
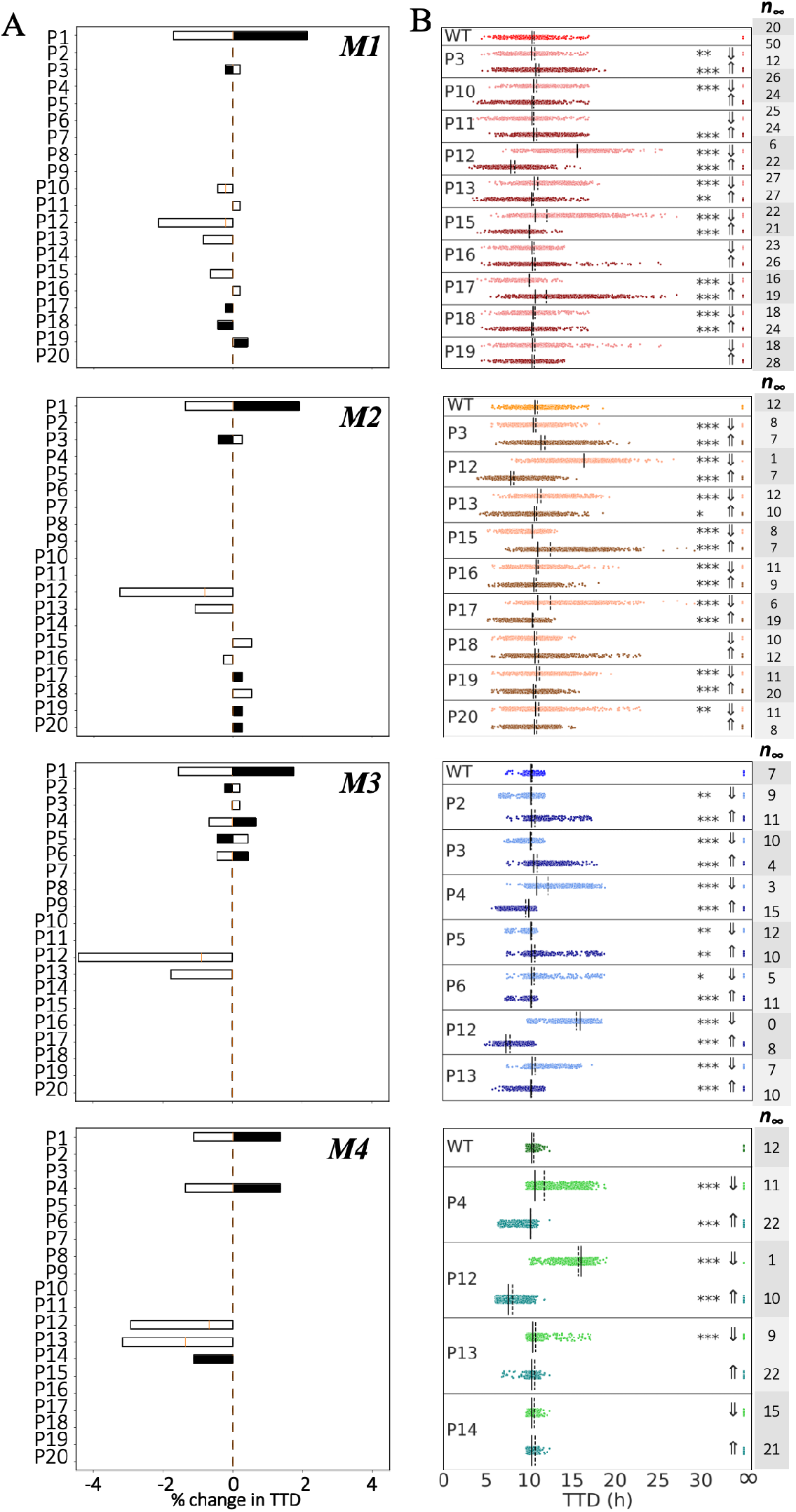
Sensitivity analyses and model-predicted rate constant targets for each mode of execution. (A) Changes in TTD for “representative” parameter sets of each mode. Black shaded regions signify decreases in rate constant values; white shaded regions signify increases. (B) Decreases (⇓; 10-fold) and increases (⇑; 10-fold) of potential targets identified in *A* for all parameter sets for each mode. The number of parameter sets that do not result in cell death (*n*_∞_) are included. Solid black lines = medians, dashed black lines = means; * p < 0.05, ** p < 0.01, *** p < 0.001 (Mood’s median test).

Across the four modes of execution, we see three common protein modulators of necroptosis sensitivity: TNF, TNFR, and MLKL (Fig. 4). These are not unexpected (and, hence, not novel targets), since these proteins are well-known master regulators of TNF-induced necroptosis.^56,57^ More interestingly, for the rate constants, we see three common modulators across the four modes (Fig. 5 and Supplementary Fig. S5) corresponding to the association of TNF to TNFR (P1), ubiquitination of RIP1 by cIAP in complex I (P12), and association of LUBAC (P13) to complex I (see Fig. 1, *pink*). The former is not unexpected, given that TNF is the death-inducing stimulus driving necroptosis. However, the latter two are not intuitively obvious and, hence, are potential global targets predicted by our model. Specifically, for all four modes, we see that increasing the values of these two rate constants (P12 and P13) leads to a significant decrease in TTD (i.e., increased sensitivity to necroptosis), and vice versa. Note that the analyses based on the representative parameter set (Fig. 5A) show only that TTD decreases when these two rate constant values are increased. However, by repeating the analyses over all parameter sets associated with each mode (Fig. 5B), we confirm that TTD also increases (i.e., sensitivity to necroptosis decreases) when the rate constant values are decreased.

### Sensitivities to initial protein levels and rate constant values reveal execution mode-dependent targets for modulating time-to-death

We have shown that the four modes of necroptosis execution (Fig. 3A) exhibit differences in variability in TTD (Fig. 3B), rate parameter values (Fig. 3C), and responses to A20 and CYLD KOs (Fig. 3D). This suggests that, in addition to the global modulators identified above (TNF, TNFR, MLKL, P1, P12, P13; Figs. 4 and 5), each mode also has a unique set of factors that drive response. For mode 1, these include proteins, i.e., A20, cIAP, and CYLD (Fig. 4–*top row*), and rate constants (P10, P11, P15–P19; Fig. 5–*top row* and Supplementary Fig. S5) associated with RIP1 ubiquitination regulation in complex I (see Fig. 1, *orange*). The sensitivities to A20 and CYLD are consistent with the results from *in silico* KO experiments (Fig. 3D). Intuitively, we can understand these sensitivities as due to competitive binding between A20 and CYLD to complex I coupled with differences in the rate constants for RIP1 deubiquitination by A20 (P19) and CYLD (P20; see Fig. 3C). In other words, increasing the amount of A20 leads to increased amounts of A20-bound complex I (and vice versa). Since the rate constant for RIP1 deubiquitination in mode 1 by A20 is much larger than for CYLD (Fig. 3C), this results in a significant decrease in TTD (i.e., increase in sensitivity to necroptosis). Conversely, increasing the amount of CYLD leads to more CYLD-bound complex I (and vice versa). Since CYLD is less efficient at deubiquitinating RIP1, this results in a much lower overall rate of RIP1 deubiquitination and a significant increase in TTD (decrease in sensitivity to necroptosis). Sensitivities to rate constants associated with these processes (P10, P11, P15–P19) can be explained similarly.

As in mode 1, potential targets in mode 2 include proteins, i.e., A20, CYLD, and LUBAC (Fig. 4A, *second row*), and rate constants (P15–P20; Fig. 5–*second row* and Supplementary Fig. S5) associated with RIP1 ubiquitination regulation. The sensitivities to A20 and CYLD, however, are reversed in their effects on TTD as compared to mode 1, i.e., increasing A20 increases TTD, while increasing CYLD decreases TTD. Again, these results are consistent with *in silico* KO experiments (Fig. 3D) and can be understood in terms of competitive binding between A20 and CYLD to complex I and differences in rate constants for RIP1 deubiquitination by A20 and CYLD (Fig. 3C). Also note that TTD in modes 1 and 2 are sensitive to the rate constant for TNF degradation (P3; Fig. 5–*top and second rows*), which is not unexpected since TNF is the stimulus driving necroptosis.

For mode 3, potential targets are associated with formation of the necrosome from complex IIb, which immediately precedes necroptosis execution (see Fig. 1, *blue*). Specifically, we see sensitivities to proteins C8, RIP1, and TRADD (Fig. 4–*third row*), the latter two of which are key components of complex II, and rate constants (P2–P6; Fig. 5–*third row* and Supplementary Fig. S5) for reactions upstream of complex II that include the association of RIP1 and TRADD to complex I. Intuitively, the comparatively small value of the rate constant in mode 3 for degradation of C8a:FLIP-bound complex IIb (P34; see Fig. 3C) is what ultimately drives these sensitivities. Modifying rates of reactions that contribute to complex II formation and/or the rate of binding of C8i to complex II, alters the balance between the rates of necrosome formation and degradation of complex IIb that prevents necroptosis, thus affecting TTD. Also note, in contrast to modes 1 and 2, the lack of sensitivity in mode 3 to variations in the initial concentrations of A20 and CYLD. This is because, in this mode, A20 and CYLD are effectively indistinguishable enzymes, i.e., rate constants for binding and unbinding from complex I (P15–P18) and RIP1 deubiquitination (P19 and P20) are virtually identical for both (Fig. 3C and Supplementary Fig. S5). Thus, varying the concentration of one is effectively equivalent to varying the concentration of the other by the same amount.

In mode 4, we see the same sensitivities as in mode 3 to varying concentrations of C8, RIP1, and TRADD (Fig. 4– *bottom row*) and the rate constant for association of TRADD to complex I (P4; Fig. 5–*bottom row* and Supplementary Fig. S5). These sensitivities can be understood in the same way as in mode 3, in terms of the balance between necrosome formation and complex IIb degradation. However, we see an additional sensitivity in mode 4 to the initial concentration of LUBAC (Fig. 4–*bottom row*). Interestingly, for the representative parameter set, this is evident for both increases and decreases in LUBAC concentration (Fig. 4A–*bottom row*), but when all parameter sets are considered is only statistically significant for the KD experiments (Fig. 5A– *bottom row*). Note also that the representative parameter set shows a sensitivity to the dissociation rate of LUBAC from complex I (P14; Fig. 5A– *bottom row*) but the effect is not statistically significant when all parameter sets are considered (Fig. 5B–*bottom row*). Furthermore, despite the results of *in silico* KO experiments that show RIP1 deubiquitination in mode 4 is driven exclusively by CYLD (Fig. 3D), we do not see a sensitivity in TTD to variations in CYLD concentration, even for a 70% KD (Fig. 4–*bottom row*). We can explain both this result and the one-way sensitivity to variations in LUBAC as due to a severely dysfunctional A20 in mode 4, evident in exceedingly small rate constants for A20 binding to complex I (P15) and subsequent RIP1 deubiquitination (P19), coupled with a comparatively large rate constant for C8 activation (P27) and small rate constant for C8 inactivation (P28; Fig. 3C). Essentially, A20 does not compete with CYLD for binding to complex I (P15 ≪ P16), and since CYLD is in great excess relative to complex I (Supplementary Fig. S6A), varying CYLD concentration has little to no effect on TTD except for very large reductions, such as a KO (Fig. 3D and Supplementary Fig. S6B). Moreover, the exceedingly fast rate of C8 activation (P28/P27 ≪ 1) leads to a rapid accumulation of active C8a:FLIP heterodimer, which inhibits necroptosis by binding and degrading complex IIb. This essentially sets a “speed limit” on the rate of pMLKL production, i.e., any increase in complex I concentration due to an increase in the concentration of LUBAC, which would be expected to decrease TTD because of the large excess of CYLD, is counteracted by the increased concentration of C8a:FLIP. However, decreasing complex I concentration by knocking down LUBAC would still be expected to increase TTD, as confirmed by our results.

## DISCUSSION

A recent review of TNF-induced necroptosis^56^ described signaling along the RIP1-RIP3-MLKL axis in terms of at least three major compartmentalization events: TNFR internalization in complex I, multiprotein assembly of complexes IIa and IIb, and necrosome formation leading to translocation of pMLKL to the membrane. Importantly, the authors emphasized that cues and regulation mechanisms underlying these compartmentalization events are poorly understood and proposed that a network of modulators surrounds the necroptotic signaling core,^58–60^ tuned in a context-, cell type-, and species-dependent manner. The results presented here are entirely consistent with this view, i.e., a detailed kinetic model comprising core and complementary necroptotic signaling proteins and associated rate constants (Table 1 and Fig. 1), calibrated to experimental data (Fig. 2A–C), can produce cell-death dynamics via distinct execution modes (Fig. 3A,B), distinguished by variations in rate constants (Fig. 3C) and the roles of A20 and CYLD in RIP1 ubiquitination regulation (Table 2 and Fig. 3D). Moreover, model sensitivity analyses based on TTD (Fig. 2D) revealed global and mode-specific modulators of necroptosis sensitivity for each mode (Figs. 4 and 5). Global modulators include known effectors, such as TNF, TNFR, MLKL, and rate constants associated with these proteins, as well as two unexpected modulators: the rate constant for RIP1 ubiquitination by cIAP in complex I (P12) and the binding rate constant for LUBAC to complex I (P13). Mode-specific modulators include, for modes 1 and 2, proteins and rate constants involved in RIP1 ubiquitination regulation (A20, cIAP, CYLD, LUBAC, P10, P11, P15–P20) and, for modes 3 and 4, factors regulating the balance between complex IIb degradation and necrosome formation (C8, LUBAC, RIP1, TRADD, P2–P6, P14, P27, P28).

In addition, numerous published experimental studies have shown that RIP1 deubiquitination in complex I is driven by A20, CYLD, or both, depending on cell type. For example, Wertz et al.^22^ showed that A20 can deubiquitinate RIP1 in human embryonic kidney (HEK) cells and mouse embryonic fibroblasts (MEFs). In contrast, Feoktistova et al.^61^ reported that deletion of A20 in human T lymphocyte (HTL) cells has no effect on necroptosis sensitivity. Moreover, Moquin et al.^12^ reported that RIP1 deubiquitination in MEFs is mediated by CYLD, but proposed it occurs in the necrosome rather than complex I, since KD of CYLD had no effect on RIP1 deubiquitination. Vanlangenakker et al.^26^ showed in mouse fibrosarcoma (MFS) cells that RIP1 can be deubiquitinated by both A20 and CYLD but, while inhibition of CYLD protects cells from necroptosis, inhibiting A20, counterintuitively, increases sensitivity to necroptosis. They also observed no effect on necroptosis after KD of TRADD. Hitomi et al.^10^ showed that increased CYLD expression reduces necroptosis in HTL cells. Similarly, Liu et al.^62^ showed in hippocampal neurons (HCNs) that KD of CYLD blocks necroptosis and Wright et al.^20^ showed that CYLD deubiquitinates RIP1 in human cervical adenocarcinoma (HCAC) cells.

To reconcile these contrasting reports, we have associated with each experimental study one or more modes of necroptosis execution identified via our model analysis (Table 3). Specifically, the report by Wertz et al.^22^ that A20 deubiquitinates RIP1 in HEK cells and MEFs implies that knocking down A20 would lead to an increase in TTD, i.e., a decrease in sensitivity to necroptosis, which is consistent with mode 1 (Fig. 3D). Conversely, the reports by Hitomi et al.^10^, Liu et al.^62^, and Wright et al.^20^ all suggest that knocking down CYLD would increase TTD, which could be explained by either modes 2 or 4 (Fig. 3D). The report by Vanlangenakker et al.^26^ also suggests that knocking down CYLD would increase TTD but, importantly, includes additional data that excludes mode 4 as a possibility, i.e., KD of A20, counterintuitively, increases sensitivity to necroptosis and TRADD KD has no effect, which are only consistent with mode 2 (Fig. 3D and Fig. 4–*second and bottom rows*). The observation by Feoktistova et al.^61^ that deletion of A20 has no effect on necroptosis sensitivity in HCAC cells is intriguing because it is consistent with both modes 3 and 4 (Fig. 3D) and they used the same cell line (HeLa) as Wright et al.^20^, who’s observations are consistent with modes 2 and 4 (as mentioned above). This could indicate that HCAC cells (or HeLa cells, specifically) operate via mode 4, since both studies are consistent with this mode, or that the cells in these experiments are operating via different modes of necroptosis execution due to differences in context, i.e., genetic or epigenetic variations between samples or differences in experimental conditions between laboratories. Finally, the report by Moquin et al.^12^ is particularly interesting because their observation that CYLD binds to complex I but RIP1 ubiquitination is not affected in CYLD-deficient MEFs led them to conclude that RIP1 ubiquitination is regulated by CYLD in the necrosome, rather than complex I. However, our analysis shows these observations are consistent with mode 4, in which TTD increases for CYLD KO (Fig. 3D) but there is no effect on TTD for CYLD KD < 90% (Fig. 4B–*bottom row* and Supplementary Fig. 6B). Thus, the results of our *in silico* analyses, based on different parameterizations of a consensus model of necroptosis, can explain a variety of incommensurate and counterintuitive experimental observations in the literature and provide an alternate explanation for a result that is seemingly inconsistent with prior studies.

**Table 3.** Multiple experimental studies of necroptosis in the literature can be associated with different model-predicted modes of execution. In the seconds column, the specific cell line used (if applicable) is included in parentheses. HCAC: human cervical adenocarcinoma; HCN: hippocampal neuron; HEK: human embryonic kidney; HTL: human T lymphocyte; MEF: mouse embryonic fibroblast; MFS: mouse fibrosarcoma. ⇓: decrease; ⇑: increase; ⇔: no change.

| Reference | Cell type | Quote(s) from article | Interpretation | Possible execution mode(s) |
| --- | --- | --- | --- | --- |
| Feoktistova et al. (2020) | HCAC (HeLa) | "[T]he deletion of A20 in HeLa or HaCaT cells had no effect on the TNF-mediated cell death sensitivity" | A20 ↓ TTD ⇔ | M3, M4 |
| Hitomi et al. (2008) | HTL (Jurkat) | "[I]nhibition of CYLD expression in Jurkat cells also attenuated necroptosis" | CYLD ↓ TTD ↑ | M2, M4 |
| Liu et al. (2014) | HCN (HT-22) | "RIP1 and its deubiquitinase CYLD are required for TNF-induced necrosis of HT-22 cells" | CYLD ↓ TTD ↑ | M2, M4 |
| Moquin et al. (2013) | MEF | "CYLD regulates RIP1 ubiquitination in the TNFα-induced necrosome, but not in the TNFR-1 signaling complex"<br>"Although CYLD was recruited to TNFR-1 in a ligand-dependent manner, RIP1 ubiquitination was not affected in CYLD<sup>-/-</sup> MEFs" | CYLD ↓ TTD ↑ | M4<br>(M2 excluded; see text) |
| Vanlangenakker et al. (2011) | MFS (L929) | "[W]e and others previously showed that CYLD repression protects L929 cells from TNF-induced necroptosis"<br>"[W]e were surprised to find that A20 depletion had an opposite effect and greatly sensitized the cells to death"<br>"[W]e found that TRADD depletion in L929 cells did not affect TNF-induced necroptosis" | CYLD ↓ TTD ↑<br>A20 ↓ TTD ↓<br>TRADD ↓ TTD ⇔ | M2 |
| Wertz et al. (2004) | HEK (HEK293T) | "Co-transfection of wild-type A20 de-ubiquitinates RIP in HEK293T cells." | A20 ↓ TTD ↑ | M1 |
| Wertz et al. (2004) | MEF | "However, in the absence of A20, RIP1 will neither be de-ubiquitinated nor targeted for proteasomal degradation. Indeed, RIP recruited to activated TNFR1 remained hyperubiquitinated and was stabilized in A20<sup>-/-</sup> MEFs" | A20 ↓ TTD ↑ | M1 |
| Wright et al. (2007) | HCAC (HeLa) | "RIP1 ubiquitination [was] inhibited by wild-type (Wt) CYLD but not a catalytically inactive CYLD mutant (Mut)" | CYLD ↓ TTD ↑ | M2, M4 |

Since evading apoptosis is a hallmark of cancer,^5–7^ inducing necroptosis is currently being explored as a potential anticancer treatment.^36,53,63^ Moreover, inhibiting necroptosis is crucial for treating a variety of inflammatory diseases, including cardiovascular, liver, and neurodegenerative diseases.^4,13^ Thus, improving our understanding of the molecular pathways that drive necroptosis is critical for identifying novel therapeutic targets against these deadly diseases. The detailed kinetic model of TNF-induced necroptosis proposed in this work represents the first successful attempt to describe contrasting, and sometimes counterintuitive, context-, cell type-, and species-dependent responses to cell-death cues using a consensus set of biochemical interactions deduced from decades of experimental work. This is a significant contribution that advances our knowledge of necroptosis and also provides a foundation for future *in silico*-guided drug discovery efforts. For example, the model can be expanded to include additional proteins and small molecules known to play a role in necroptosis^64,65^ (e.g., ADAM17, CHIP, TAK1, nerostatins), additional necroptosis-associated receptors^9^ (e.g., TNFR2, CD95, Toll-like receptors) and ligands^66–68^ (e.g., LPS, FasL, TRAIL), both forms of cIAP^69^ (i.e., cIAP1 and cIAP2), assembly of the LUBAC trimer complex,^70^ different RIP1 ubiquitin chains^56^ (i.e., M1, K48, K63), and additional biochemical events involved in the activation of C8^71^ (e.g., binding of pro-C8 to FADD, followed by oligomerization and cleavage) and formation of the necrosome^72^ (e.g., RIP3 phosphorylation by CK1 family kinases). The model can also be extended to include downstream events involved in MLKL-mediated permeabilization of the plasma membrane^73,74^ (e.g., Golgi-, microtubule-, and actin-dependent mechanisms), crosstalk with pro-survival^1,27^ (e.g., NF-κB) and other programmed cell death^75^ (e.g., apoptosis) pathways, and connections to the immune system^36^ (e.g., antigen-induced proliferation of T cells). Altogether, the model presented in this study is a significant step towards the construction of a comprehensive computational model of the interconnected pathways controlling cell fate decisions, which could lead to the development of novel therapies against inflammatory diseases and cancer by enabling identification of molecular targets that shift the balance of fates towards either evasion or promotion of necroptosis.

## MATERIALS AND METHODS

### Cell culture and reagents

L929 cells (NCTC clone 929, L cell, L-929, derivative of Strain L) were purchased from the American Type Culture Collection (ATCC) and cultured in Dulbecco’s Modified Eagle Medium (DMEM; Corning) supplemented with 10% fetal bovine serum (FBS; Omega Scientific), 1% L–Glutamine, and 1% penicillin/streptomycin (Thermo Fisher Scientific) at 5% CO2 and 37°C. Mouse recombinant TNF was purchased from R&D (Cat# 410-MT-10).

### Immunoblotting

L929 cells (2–3 × 10^6^) were grown in 10-cm dishes for 24 h followed by treatment with TNF (0.1, 1, 10, or 100 ng/ml) for 16h. Dead cells were removed by washing with ice cold phosphate-buffered saline (PBS). Remaining adherent cells were lysed using radioimmunoprecipitation assay (RIPA) buffer with 1% Triton X-100, protease, and phosphatase inhibitors. Samples were normalized for total protein concentration (Bradford assay, Bio-Rad), denaturated in 3’ sodium dodecyl sulfate (SDS) sample buffer (5 minutes at 95°C) and subjected to gel electrophoresis (4– 15% Criterion™ TGX™ Precast Midi Protein Gel, Bio-Rad) and immunoblotting (polyvinylidene difluoride Transfer Membrane, Thermo Fisher Scientific). Membranes were blocked in 5% bovine serum albumin (BSA)/tris buffered saline with Tween® 20 (TBS-T) and incubated with the following antibodies: pMLKL (1:1000, Abcam, Cat# ab196436), actin (1:3000, Santa Cruz, Cat# sc-1615), anti-rabbit (1:5000, Santa Cruz, Cat# sc-2004), anti-goat (1:3000, Santa Cruz, Cat# sc-2354). Signal was developed using chemiluminescent substrate (SuperSignal West Pico Plus, Thermo Fisher Scientific) and visualized with ChemiCoc MP imaging system (Bio-Rad).

### Determining initial protein concentrations

Expression levels for six proteins (caspase-8, FADD, unmodified MLKL, RIP3, TRADD, and TRAF2) were measured in L929 cells using absolute protein quantitation mass spectrometry. As a negative control, cells were collected in three replicate 6-well plates and cell lysates were gathered, prepped for protein precipitation, pellet, and digestion in the Vanderbilt Mass Spectrometry Research Center (MSRC) Proteomics Core Laboratory. For the other eight proteins in the model, initial concentrations were estimated from measurements reported in the literature and the human protein atlas.^76–78^ Concentrations were converted to units of molecules/cell assuming an L929 cell diameter of 15μm.^79^

### Bayesian parameter calibration

We estimated parameter values using PyDREAM,^48^ a Python implementation of the DiffeRential <u>E</u>volution <u>A</u>daptive <u>M</u>etropolis (DREAM) method.^80^ We utilized pMLKL Western blot data at the two highest TNF doses (100 and 10 ng/ml) and defined a multi-objective cost function,

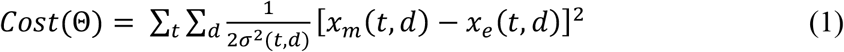

where Θ is the parameter set, *x_m_*(*t,d*) and *x_e_*(*t,d*) are model-predicted and experimentally measured pMLKL concentrations, respectively, at time *t* and TNF dose *d*, and *σ*(*t*) = 0.1·*x_e_*(*t,d*) (following previous studies^49,81,82^). Parameter sampling was performed using five Monte Carlo chains, each run for 50,000 iterations, the first 25,000 of which were considered burn-in and discarded, resulting in 125,000 parameter sets. Out of these, we extracted an ensemble of 10,628 unique parameter sets. Convergence was achieved for all chains (Supplementary Fig. S1), assessed using the Gelman-Rubin test.^83,84^ Starting positions for all PyDREAM chains were determined using particle swarm optimization^85^ (PSO): we performed 100 PSO runs, of 500 iterations each, saved the parameter sets from the last iteration of each run, and selected the five with the lowest cost function values (Eq. 1). Also, for all parameters, we set prior distributions in PyDREAM to log-normal distributions, 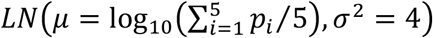, where *p*, is the value of the parameter from the *i*-th PSO run. Starting rate constant values for the PSO runs were set to physically plausible values:^86,87^ association=10^-6^ min^-1^, dissociation=10^-3^ min^-1^, ubiquitination/phosphorylation=1 min^-1^, and degradation=1 min^-1^ (see Supplementary Table S2). In all cases, simulations were performed by numerical integration of ordinary differential equations (ODEs) using LSODA,^88^ as implemented in the Python package SciPy.^89^

### Identifying modes of signal execution in a parameter set ensemble

Modes of signal execution were identified using PyDyNo, a Python-based software package for dynamical systems analysis of biochemical models with uncertain parameters.^50^ PyDyNo takes as input a model object (PySB^90^ or SBML^91,92^ formats), an input file with parameter sets, and a target species (pMLKL, in our case). ODE simulations are run^88,89^ for all parameter sets and “digitized” into a sequence of integers, termed a “dynamical signature,” based on “dominant” subnetworks of reactions identified at each time point. Basically, the algorithm identifies, at every time point, the subnetwork of reactions that contribute most to either the production or consumption (depending on user preference; production, in our case) of the target species and assigns to each identified subnetwork an integer index. Each time point is thus associated with an integer index and the entire simulated time course with a sequence of integers, i.e., the dynamical signature. We refer the reader to the original work^50^ for further details on how PyDyNo identifies dominant subnetworks from ODE simulations of biochemical models. We repeated this procedure for all 10,628 unique parameter sets obtained from PyDREAM, with all simulations run at the highest TNF dose (100 ng/ml) for 16h simulated time, in line with experimental data (Fig. 2A). Dynamical signatures were clustered using a spectral clustering method^93^ with the longest common subsequence^51^ (LCS) as the distance metric. The optimal number of clusters, i.e., modes of execution, was determined using a silhouette score^94^ for cluster sizes between 2 and 20 (Supplementary Fig. S3). For each mode, a “representative” dynamical signature was defined as the one with the minimal sum of distances to all other signatures^95^ (i.e., the medoid).

### Sensitivity analyses for initial protein concentrations and rate constants

We used a sensitivity analysis tool^96^ available in PySB^90^ to quantify changes in TTD, defined as the time at which pMLKL reaches a pre-defined threshold (Fig. 2D), due to changes in both initial protein concentrations and rate constants. Briefly, the sensitivity analysis tool varies pairs of protein concentrations or rate constants over a range of values relative to a reference set (in this case, [−20%, …, −2%, 0%, 2%, …, 20%]) and calculates the resulting changes in TTD. For each protein or rate constant, a “single-parameter sensitivity multiset^96^” is then obtained, which summarizes the range of changes in TTD due to the changes in protein or rate constant values and can be visualized as a boxplot (Figs. 4A and 5A). Reference rate constants are those associated with the representative dynamical signatures obtained for each mode from PyDyNo (see previous subsection). For protein concentration sensitivities, reference concentrations are those obtained from mass spectrometry (Fig. 2B) and the literature or human protein atlas^76–78^ (Supplementary Table S1) and all simulations were performed using the reference rate constant values. Note that we defined a hard threshold of 2,772 pMLKL molecules to define TTD, which is half the amount measured by mass spectrometry (Fig. 2B). We chose this, rather than, e.g., the half-maximal amount of pMLKL, to prevent any bias (i.e., changes in the threshold) when varying the initial amount of MLKL. This choice is consistent with experimental evidence that plasma membrane damage accumulates until a threshold is reached, triggering cell death.^74^ Results of the sensitivity analyses above, which used reference rate constant values, were then validated by performing, over the full set of rate constant values for each mode, *in silico* KD (70%) and OE (10-fold) experiments for protein concentrations and ± 10-fold variations for the rate constants (Figs. 4B and 5B). This was critical for identifying results that were specific only to the reference parameter set and, hence, could be discounted from our analyses.

## Data and computer code availability

All Western blot data, mass spectrometry data, and Python code used in this study, including the PySB encoding of the Necroptosis Execution Reaction Model (NERMv1.0), are available at https://github.com/LoLab-VU/NERM.git.

## Acknowledgments

This work is dedicated to the memory of our friend and colleague Melaine N. Sebastian. We thank Sam Beik, Sarah Maddox Groves, Corey Hayford, Michael Irvin, Alex Lubbock, Tolu Omokehinde, Oscar Ortega, James Pino, and Vito Quaranta for useful discussions regarding this work and Hayes McDonald of the Vanderbilt MSRC Proteomics Core Laboratory for help in obtaining the mass spectrometry data. G.V.I. also thanks Beth Bowman, Don Brunson, Roger Chalkley, Christina Keeton, Linda Sealy, and Patricia Mueller of Vanderbilt University, and Michael Aldarando-Jeffries, Arlene Olivierre, and Natalia Toro of the University of Central Florida for continued support.

## Author contributions

Conceptualization (AH, CFL, GVI)

Methodology (CFL, GVI, LAH)

Resources (AH, CFL)

Software (CFL, GVI)

Formal Analysis (CFL, GVI, LAH)

Investigation (AH, CFL, GVI, LAH, MOM)

Writing – Original Draft (CFL, GVI, LAH)

Writing – Review and Editing (AH, CFL, GVI, LAH, MOM)

Visualization (CFL, GVI, LAH)

Project Administration (AH, CFL, LAH)

Funding Acquisition (AH, CFL, GVI)

## REFERENCES

1 Vanden Berghe T, Kaiser WJ, Bertrand MJM, Vandenabeele P. Molecular crosstalk between apoptosis, necroptosis, and survival signaling. Mol Cell Oncol 2015; 2: e975093.

2 Degterev A, Huang Z, Boyce M, Li Y, Jagtap P, Mizushima N et al. Chemical inhibitor of nonapoptotic cell death with therapeutic potential for ischemic brain injury. Nat Chem Biol 2005; 1: 112–119.

3 Aldridge BB, Saez-Rodriguez J, Muhlich JL, Sorger PK, Lauffenburger DA. Fuzzy logic analysis of kinase pathway crosstalk in TNF/EGF/insulin-induced signaling. PLoS Comput Biol 2009; 5: e1000340.

4 Vanlangenakker N, Vanden Berghe T, Vandenabeele P. Many stimuli pull the necrotic trigger, an overview. Cell Death Differ 2012; 19: 75–86.

5 Hanahan D, Weinberg RA. The hallmarks of cancer. Cell 2000; 100: 57–70.

6 Hanahan D, Weinberg RA. Hallmarks of cancer: the next generation. Cell 2011; 144: 646–74.

7 Hanahan D. Hallmarks of cancer: new dimensions. Cancer Discov 2022; 12: 31–46.

8 Chan K, Saltelli A, Tarantola S. Sensitivity analysis of model output: Variance-based methods make the difference. In: Winter Simulation Conference Proceedings. 1997, pp 261–268.

9 Zhou W, Yuan J. Necroptosis in health and diseases. Semin Cell Dev Biol 2014; 35: 14–23.

10 Hitomi J, Christofferson DE, Ng A, Yao J, Degterev A, Xavier RJ et al. Identification of a molecular signaling network that regulates a cellular necrotic cell death pathway. Cell 2008; 135: 1311–1323.

11 Vanlangenakker N, Vanden Berghe T, Bogaert P, Laukens B, Zobel K, Deshayes K et al. cIAP1 and TAK1 protect cells from TNF-induced necrosis by preventing RIP1/RIP3-dependent reactive oxygen species production. Cell Death Differ 2011; 18: 656–665.

12 Moquin DM, McQuade T, Chan FK-MM. CYLD deubiquitinates RIP1 in the TNFα-induced necrosome to facilitate kinase activation and programmed necrosis. PLoS One 2013; 8: e76841.

13 Choi ME, Price DR, Ryter SW, Choi AMK. Necroptosis: A crucial pathogenic mediator of human disease. JCI Insight 2019; 4: e128834.

14 Wartz IE, O’Rourke KM, Zhou H, Eby M, Aravind L, Seshagiri S et al. De-ubiquitination and ubiquitin ligase domains of A20 downregulate NF-κB signalling. Nature 2004; 430: 694–699.

15 Sun SC. A20 restricts inflammation via ubiquitin binding. Nat Immunol 2020; 21: 362–364.

16 Lork M, Verhelst K, Beyaert R. CYLD, A20 and OTULIN deubiquitinases in NF-κB signaling and cell death: So similar, yet so different. Cell Death Differ 2017; 24: 1172–1183.

17 Simonson SJS, Wu ZH, Miyamoto S. CYLD: A DUB with many talents. Dev Cell 2007; 13: 601–603.

18 Vandenabeele P, Galluzzi L, Vanden Berghe T, Kroemer G. Molecular mechanisms of necroptosis: an ordered cellular explosion. Nat Rev Mol Cell Biol 2010; 11: 700–714.

19 Kovalenko A, Chable-Bessia C, Cantarella G, Israël A, Wallach D, Courtois G. The tumour suppressor CYLD negatively regulates NF-κB signalling by deubiquitination. Nature 2003; 424: 801–805.

20 Wright A, Reiley WW, Chang M, Jin W, Lee AJ, Zhang M et al. Regulation of early wave of germ cell apoptosis and spermatogenesis by deubiquitinating enzyme CYLD. Dev Cell 2007; 13: 705–716.

21 Gurung P, Man SM, Kanneganti T-D. A20 is a regulator of necroptosis. Nat Immunol 2015; 16: 596–597.

22 Wertz IE, O’Rourke KM, Zhou H, Eby M, Aravind L, Seshagiri S et al. De-ubiquitination and ubiquitin ligase domains of A20 downregulate NF-κB signalling. Nature 2004; 430: 694–699.

23 Lu TT, Onizawa M, Hammer GE, Turer EE, Yin Q, Damko E et al. Dimerization and ubiquitin mediated recruitment of A20, a complex deubiquitinating enzyme. Immunity 2013; 38: 896–905.

24 Dondelinger Y, Darding M, Bertrand MJM, Walczak H. Poly-ubiquitination in TNFR1-mediated necroptosis. Cell Mol Life Sci 2016; 73: 2165–2176.

25 Draber P, Kupka S, Reichert M, Draberova H, Lafont E, de Miguel D et al. LUBAC-recruited CYLD and A20 regulate gene activation and cell death by exerting opposing effects on linear ubiquitin in signaling complexes. Cell Rep 2015; 13: 2258–2272.

26 Vanlangenakker N, Bertrand MJM, Bogaert P, Vandenabeele P, Vanden Berghe T. TNF-induced necroptosis in L929 cells is tightly regulated by multiple TNFR1 complex I and II members. Cell Death Dis 2011; 2: e230.

27 Metzig MO, Tang Y, Mitchell S, Taylor B, Foreman R, Wollman R et al. An incoherent feedforward loop interprets NFκB/RelA dynamics to determine TNF-induced necroptosis decisions. Mol Syst Biol 2020; 16: e9677.

28 Li M, Beg AA. Induction of necrotic-like cell death by tumor necrosis factor alpha and caspase inhibitors: novel mechanism for killing virus-infected cells. J Virol 2000; 74: 7470–7477.

29 Wallach D. Preparations of lymphotoxin induce resistance to their own cytotoxic effect. J Immunol 1984; 132: 2464–2469.

30 Wajant H, Siegmund D. TNFR1 and TNFR2 in the control of the life and death balance of macrophages. Front Cell Dev Biol 2019; 7: 91.

31 Van Antwerp DJ, Martin SJ, Kafri T, Green DR, Verma IM. Suppression of TNF-α-induced apoptosis by NF-κB. Science (80-) 1996; 274: 787–789.

32 Pobezinskaya YL, Liu Z. The role of TRADD in death receptor signaling. Cell Cycle 2012; 11: 871–876.

33 Zheng L, Bidere N, Staudt D, Cubre A, Orenstein J, Chan FK et al. Competitive control of independent programs of tumor necrosis factor receptor-induced cell death by TRADD and RIP1. Mol Cell Biol 2006; 26: 3505–3513.

34 Liu X, Shi F, Li Y, Yu X, Peng S, Li W et al. Post-translational modifications as key regulators of TNF-induced necroptosis. Cell Death Dis 2016; 7: e2293.

35 Etemadi N, Chopin M, Anderton H, Tanzer MC, Rickard JA, Abeysekera W et al. TRAF2 regulates TNF and NF-κB signalling to suppress apoptosis and skin inflammation independently of sphingosine kinase. Elife 2015; 4: e10592.

36 Gong Y, Fan Z, Luo G, Yang C, Huang Q, Fan K et al. The role of necroptosis in cancer biology and therapy. Mol Cancer 2019; 18: 100.

37 Micheau O, Tschopp J. Induction of TNF receptor I-mediated apoptosis via two sequential signaling complexes. Cell 2003; 114: 181–190.

38 Micheau O, Thome M, Schneider P, Holler N, Tschopp J, Nicholson DW et al. The long form of FLIP Is an activator of caspase-8 at the Fas death-inducing signaling complex. J Biol Chem 2002; 277: 45162–45171.

39 Tsuchiya Y, Nakabayashi O, Nakano H. FLIP the switch: regulation of apoptosis and necroptosis by cFLIP. Int J Mol Sci 2015; 16: 30321–30341.

40 McIlwain DR, Berger T, Mak TW. Caspase functions in cell death and disease. Cold Spring Harb Perspect Biol 2013; 5: a008656.

41 Feoktistova M, Geserick P, Kellert B, Dimitrova DP, Langlais C, Hupe M et al. CIAPs block Ripoptosome formation, a RIP1/caspase-8 containing intracellular cell death complex differentially regulated by cFLIP isoforms. Mol Cell 2011; 43: 449–463.

42 Moriwaki K, Chan FK-M. RIP3: a molecular switch for necrosis and inflammation. Genes Dev 2013; 27: 1640–1649.

43 Declercq W, Vanden Berghe T, Vandenabeele P. RIP kinases at the crossroads of cell death and survival. Cell 2009; 138: 229–232.

44 Sun L, Wang H, Wang Z, He S, Chen S, Liao D et al. Mixed lineage kinase domain-like protein mediates necrosis signaling downstream of RIP3 kinase. Cell 2012; 148: 213–227.

45 Cho YS, Challa S, Moquin D, Genga R, Ray TD, Guildford M et al. Phosphorylation-driven assembly of the RIP1-RIP3 complex regulates programmed necrosis and virus-induced inflammation. Cell 2009; 137: 1112–1123.

46 Ronan T, Qi Z, Naegle KM, Jain AK, Murty MN, Flynn PJ et al. Avoiding common pitfalls when clustering biological data. Sci Signal 2016; 9: re6.

47 Pasparakis M, Vandenabeele P. Necroptosis and its role in inflammation. Nature 2015; 517: 311–320.

48 Shockley EM, Vrugt JA, Lopez CF, Valencia A. PyDREAM: high-dimensional parameter inference for biological models in python. Bioinformatics 2018; 34: 695–697.

49 Eydgahi H, Chen WW, Muhlich JL, Vitkup D, Tsitsiklis JN, Sorger PK. Properties of cell death models calibrated and compared using Bayesian approaches. Mol Syst Biol 2013; 9: 644.

50 Ortega OO, Wilson BA, Pino JC, Irvin MW, Ildefonso GV, Garbett SP et al. Probability-based mechanisms in biological networks with parameter uncertainty. bioRxiv 2021. doi:10.1101/2021.01.26.428266.

51 Studer M, Ritschard G. What matters in differences between life trajectories: a comparative review of sequence dissimilarity measures. J R Stat Soc Ser A Stat Soc 2016; 179: 481–511.

52 Rokach L, Maimon O. Clustering methods. In: Maimon O, Rokach L (eds). Data Mining and Knowledge Discovery Handbook. Springer-Verlag, 2005, pp 321–349.

53 Wu Y, Dong G, Sheng C. Targeting necroptosis in anticancer therapy: mechanisms and modulators. Acta Pharm Sin B 2020; 10: 1601–1618.

54 Taxman DJ, Livingstone LR, Zhang J, Conti BJ, Iocca HA, Williams KL et al. Criteria for effective design, construction, and gene knockdown by shRNA vectors. BMC Biotechnol 2006; 6: 7.

55 Moriya H. Quantitative nature of overexpression experiments. Mol Biol Cell 2015; 26: 3932.

56 Samson AL, Garnish SE, Hildebrand JM, Murphy JM. Location, location, location: A compartmentalized view of TNF-induced necroptotic signaling. Sci Signal 2021; 14: 6178.

57 Vercammen D, Vandenabeele P, Beyaert R, Declercq W, Fiers W. Tumour necrosis factor-induced necrosis versus anti-Fas-induced apoptosis in L929 cells. Cytokine 1997; 9: 801–808.

58 Murphy JM, Czabotar PE, Hildebrand JM, Lucet IS, Zhang J-G, Alvarez-Diaz S et al. The pseudokinase MLKL mediates necroptosis via a molecular switch mechanism. Immunity 2013; 39: 443–453.

59 Najafov A, Mookhtiar AK, Luu HS, Ordureau A, Pan H, Amin PP et al. TAM kinases promote necroptosis by regulating oligomerization of MLKL. Mol Cell 2019; 75: 457–468.e4.

60 Sai K, Parsons C, House JS, Kathariou S, Ninomiya-Tsuji J. Necroptosis mediators RIPK3 and MLKL suppress intracellular Listeria replication independently of host cell killing. J Cell Biol 2019; 218: 1994–2005.

61 Feoktistova M, Makarov R, Brenji S, Schneider AT, Hooiveld GJ, Luedde T et al. A20 promotes ripoptosome formation and TNF-induced apoptosis via cIAPs regulation and NIK stabilization in keratinocytes. Cells 2020; 9: 351.

62 Liu S, Wang X, Li Y, Xu L, Yu X, Ge L et al. Necroptosis mediates TNF-induced toxicity of hippocampal neurons. Biomed Res Int 2014; 2014: 290182.

63 Meng M-B, Wang H-H, Cui Y-L, Wu Z-Q, Shi Y-Y, Zaorsky NG et al. Necroptosis in tumorigenesis, activation of anti-tumor immunity, and cancer therapy. Oncotarget 2016; 7: 57391–57413.

64 Chen J, Kos R, Garssen J, Redegeld F. Molecular insights into the mechanism of necroptosis: the necrosome as a potential therapeutic target. Cells 2019; 8: 1486.

65 Bolik J, Krause F, Stevanovic M, Gandraß M, Thomsen I, Schacht SS et al. Inhibition of ADAM17 impairs endothelial cell necroptosis and blocks metastasis. J Exp Med 2021; 219: 219.

66 Strasser A, Jost PJ, Nagata S. The many roles of FAS receptor signaling in the immune system. Immunity 2009; 30: 180–192.

67 Kearney CJ, Cullen SP, Tynan GA, Henry CM, Clancy D, Lavelle EC et al. Necroptosis suppresses inflammation via termination of TNF-or LPS-induced cytokine and chemokine production. Cell Death Differ 2015; 22: 1313–1327.

68 Füllsack S, Rosenthal A, Wajant H, Siegmund D. Redundant and receptor-specific activities of TRADD, RIPK1 and FADD in death receptor signaling. Cell Death Dis 2019; 10: 122.

69 McComb S, Cheung HH, Korneluk RG, Wang S, Krishnan L, Sad S. cIAP1 and cIAP2 limit macrophage necroptosis by inhibiting Rip1 and Rip3 activation. Cell Death Differ 2012; 19: 1791–1801.

70 Haas TL, Emmerich CH, Gerlach B, Schmukle AC, Cordier SM, Rieser E et al. Recruitment of the linear ubiquitin chain assembly complex stabilizes the TNF-R1 signaling complex and is required for TNF-mediated gene induction. Mol Cell 2009; 36: 831–844.

71 Donepudi M, Sweeney A Mac, Briand C, Grütter MG. Insights into the regulatory mechanism for caspase-8 activation. Mol Cell 2003; 11: 543–549.

72 Hanna-Addams S, Liu S, Liu H, Chen S, Wang Z. CK1α, CK1δ, and CK1ε are necrosome components which phosphorylate serine 227 of human RIPK3 to activate necroptosis. Proc Natl Acad Sci 2020; 117: 1962–1970.

73 Wang H, Sun L, Su L, Rizo J, Liu L, Wang L-F et al. Mixed lineage kinase domain-like protein MLKL causes necrotic membrane disruption upon phosphorylation by RIP3. Mol Cell 2014; 54: 133–146.

74 Samson AL, Zhang Y, Geoghegan ND, Gavin XJ, Davies KA, Mlodzianoski MJ et al. MLKL trafficking and accumulation at the plasma membrane control the kinetics and threshold for necroptosis. Nat Commun 2020; 11: 3151.

75 Ouyang L, Shi Z, Zhao S, Wang F-T, Zhou T-T, Liu B et al. Programmed cell death pathways in cancer: a review of apoptosis, autophagy and programmed necrosis. Cell Prolif 2012; 45: 487–498.

76 Hua F, Cornejo MG, Cardone MH, Stokes CL, Lauffenburger DA. Effects of Bcl-2 levels on Fas signaling-induced caspase-3 activation: Molecular genetic tests of computational model predictions. J Immunol 2005; 175: 985–995.

77 Uhlén M, Fagerberg L, Hallström BM, Lindskog C, Oksvold P, Mardinoglu A et al. Tissue-based map of the human proteome. Science (80-) 2015; 347: 1260419.

78 Uhlén M, Björling E, Agaton C, Szigyarto CAK, Amini B, Andersen E et al. A human protein atlas for normal and cancer tissues based on antibody proteomics. Mol Cell Proteomics 2005; 4: 1920–1932.

79 Milo R, Jorgensen P, Moran U, Weber G, Springer M. BioNumbers—the database of key numbers in molecular and cell biology. Nucleic Acids Res 2009; 38: D750–D753.

80 Vrugt JA, Ter Braak CJF. DREAM(D): An adaptive Markov Chain Monte Carlo simulation algorithm to solve discrete, noncontinuous, and combinatorial posterior parameter estimation problems. Hydrol Earth Syst Sci 2011; 15: 3701–3713.

81 Kochen MA, Lopez CF. A probabilistic approach to explore signal execution mechanisms with limited experimental data. Front Genet 2020; 11: 686.

82 Spencer SL, Gaudet S, Albeck JG, Burke JM, Sorger PK. Non-genetic origins of cell-to-cell variability in TRAIL-induced apoptosis. Nature 2009; 459: 428–432.

83 Vrugt JA. Markov chain Monte Carlo simulation using the DREAM software package: Theory, concepts, and MATLAB implementation. Environ Model Softw 2016; 75: 273–316.

84 Csete ME, Doyle JC. Reverse engineering of biological complexity. Science (80-.). 2002; 295: 1664–1669.

85 Marini F, Walczak B. Particle swarm optimization (PSO). A tutorial. Chemom Intell Lab Syst 2015; 149: 153–165.

86 Aldridge BB, Burke JM, Lauffenburger DA, Sorger PK. Physicochemical modelling of cell signalling pathways. Nat Cell Biol 2006; 8: 1195–1203.

87 Lawson MJ, Petzold L, Hellander A. Accuracy of the Michaelis–Menten approximation when analysing effects of molecular noise. J R Soc Interface 2015; 12: 20150054.

88 Petzold L. Automatic selection of methods for solving stiff and nonstiff systems of ordinary differential equations. SIAM J Sci Stat Comput 1983; 4: 136–148.

89 Virtanen P, Gommers R, Oliphant TE, Haberland M, Reddy T, Cournapeau D et al. SciPy 1.0: fundamental algorithms for scientific computing in Python. Nat Methods 2020; 17: 261–272.

90 Lopez CF, Muhlich JL, Bachman JA, Sorger PK. Programming biological models in Python using PySB. Mol Syst Biol 2013; 9: 646.

91 Hucka M, Finney A, Sauro HM, Bolouri H, Doyle JC, Kitano H et al. The systems biology markup language (SBML): a medium for representation and exchange of biochemical network models. Bioinformatics 2003;19: 524–531.

92 Keating SM, Waltemath D, König M, Zhang F, Dräger A, Chaouiya C et al. SBML Level 3: an extensible format for the exchange and reuse of biological models. Mol Syst Biol 2020; 16: e9110.

93 Von Luxburg U. A tutorial on spectral clustering. Stat Comput 2007 174 2007; 17: 395–416.

94 Rousseeuw PJ. Silhouettes: A graphical aid to the interpretation and validation of cluster analysis. J Comput Appl Math 1987; 20: 53–65.

95 Gabadinho A, Ritschard G, Müller NS, Studer M. Analyzing and Visualizing State Sequences in R with TraMineR. J Stat Softw 2011; 40: 1–37.

96 Harris LA, Nobile MS, Pino JC, Lubbock ALR, Besozzi D, Mauri G et al. GPU-powered model analysis with PySB/cupSODA. Bioinformatics 2017; 33: 3492–3494.

97 Feltham R, Vince JE, Lawlor KE. Caspase-8: not so silently deadly. Clin Transl Immunol 2017; 6: e124.

98 Oberst A, Dillon CP, Weinlich R, McCormick LL, Fitzgerald P, Pop C et al. Catalytic activity of the caspase-8–FLIPL complex inhibits RIPK3-dependent necrosis. Nature 2011; 471: 363–367.

99 Hsu H, Shu H-B, Pan M-G, Goeddel D V. TRADD–TRAF2 and TRADD–FADD interactions define two distinct TNF Receptor 1 signal transduction pathways. Cell 1996; 84: 299–308.

100 Lawrence CP, Chow SC. FADD deficiency sensitises Jurkat T cells to TNF-α-dependent necrosis during activation-induced cell death. FEBS Lett 2005; 579: 6465–6472.

101 Wegner KW, Saleh D, Degterev A. Complex pathologic roles of RIPK1 and RIPK3: moving beyond necroptosis. Trends Pharmacol Sci 2017; 38: 202–225.

102 Grootjans S, Vanden Berghe T, Vandenabeele P. Initiation and execution mechanisms of necroptosis: an overview. Cell Death Differ 2017; 24: 1184–1195.

103 O’Donnell MA, Ting AT. RIP1 comes back to life as a cell death regulator in TNFR1 signaling. FEBS J 2011; 278: 877–887.

104 Vandenabeele P, Declercq W, Van Herreweghe F, Berghe T Vanden. The role of the kinases RIP1 and RIP3 in TNF-induced necrosis. Sci Signal 2010; 3: re4.

105 Feng S, Yang Y, Mei Y, Ma L, Zhu D, Hoti N et al. Cleavage of RIP3 inactivates its caspase-independent apoptosis pathway by removal of kinase domain. Cell Signal 2007; 19: 2056–2067.

106 He S, Wang L, Miao L, Wang T, Du F, Zhao L et al. Receptor interacting protein kinase-3 determines cellular necrotic response to TNF-α. Cell 2009; 137: 1100–1111.

107 Carswell EA, Old LJ, Kassel RL, Green S, Fiore N, Williamson B. An endotoxin-induced serum factor that causes necrosis of tumors. Proc Natl Acad Sci U S A 1975; 72: 3666–3670.

108 Hsu H, Huang J, Shu HB, Baichwal V, Goeddel D V. TNF-dependent recruitment of the protein kinase RIP to the TNF receptor-1 signaling complex. Immunity 1996; 4: 387–396.

109 Wang CY, Mayo MW, Korneluk RG, Goeddel D V, Baldwin AS. NF-kappaB antiapoptosis: induction of TRAF1 and TRAF2 and c-IAP1 and c-IAP2 to suppress caspase-8 activation. Science 1998; 281: 1680–1683.

110 Hsu H, Shu HB, Pan MG, Goeddel D V. TRADD–TRAF2 and TRADD–FADD interactions define two distinct TNF Receptor 1 signal transduction pathways. Cell 1996; 84: 299–308.

