## Supplementary Tables and Figures for "Distinct execution modes of a biochemical necroptosis model explain cell type-specific responses and variability to cell-death cues"

**Supplementary Table S1. Initial protein levels used as input to the computational model.** Proteins shown in bold were either measured by mass spectrometry in this work or, in one case, based on applied dose (TNF). Initial amounts for the other proteins were obtained from the literature (Hua et al. 2005, Uhlén et al. 2005, Uhlén et al. 2015).

| **Protein** | **Amount (molecules)** | **Source** |
|---|---|---|
| A20 | 9,075 | Calculated |
| **C8** | 3,799 | This work |
| cIAP | 8,986 | Calculated |
| CYLD | 9,075 | Calculated |
| **FADD** | 3,109 | This work |
| FLIP | 3,900 | Calculated |
| LUBAC | 7,226 | Calculated |
| **MLKL (unmod)** | 5,544 | This work |
| RIP1 | 20,044 | Calculated |
| **RIP3** | 10,654 | This work |
| **TNF*** | 2,326 | This work |
| TNFR | 4,800 | Calculated |
| **TRADD** | 4,696 | This work |
| **TRAF2** | 11,776 | This work |

*100 ng/ml

**Supplementary Table S2: Rate parameters descriptions for all reactions in the necroptosis model.** C8i: inactive caspase-8; C8a: active caspase-8; RIP1-u: unmodified RIP1; RIP1-Ub: ubiquitinated RIP1; RIP1-dUb: deubiquitinated RIP1; RIP1-p: phosphorylated RIP1; RIP3-p: phosphorylated RIP3.

| Parameter | Reaction | Parameter | Reaction |
|---|---|---|---|
| **P1** | Association of TNF to TNFR | **P21** | Association of FADD to RIP1-dUb:TRADD in complex II |
| **P2** | Dissociation of TNF:TNFR | **P22** | Dissociation of FADD from RIP1-dUb:TRADD in complex II |
| **P3** | Degradation of TNF | **P23** | Association of C8i to FADD in complex IIa |
| **P4** | Association of TRADD to complex I | **P24** | Dissociation of C8i from complex IIa |
| **P5** | Dissociation of TRADD from complex I | **P25** | Association of FLIP to C8i in complex IIa |
| **P6** | Association of RIP1-u to complex I | **P26** | Dissociation of FLIP from C8i in complex IIa |
| **P7** | Dissociation of RIP1-u from complex I | **P27** | Activation of C8i:FLIP → C8a:FLIP heterodimer in complex IIa |
| **P8** | Association of TRAF2 to complex I | **P28** | Inactivation of C8a:FLIP → C8i:FLIP in complex IIa |
| **P9** | Dissociation of TRAF2 from complex I | **P29** | Degradation of RIP1-dUb by C8a:FLIP in complex IIa |
| **P10** | Association of cIAP to complex I | **P30** | Association RIP3 to RIP1-dUb in complex IIb |
| **P11** | Dissociation of cIAP from complex I | **P31** | Dissociation of RIP3 from RIP1-dUb in complex IIb |
| **P12** | Ubiquitination of RIP1-u by cIAP in complex I | **P32** | Association of C8a:FLIP to complex IIb |
| **P13** | Association of LUBAC to RIP1-Ub in complex I | **P33** | Dissociation of C8a:FLIP from complex IIb |
| **P14** | Dissociation of LUBAC from RIP1-Ub in complex I | **P34** | Degradation of RIP1-dUb by C8a:FLIP in complex IIb |
| **P15** | Association of A20 to RIP1-Ub in complex I | **P35** | Dissociation of RIP1-dUb:RIP3 from complex IIb |
| **P16** | Dissociation of A20 from complex I | **P36** | Phosphorylation of RIP1-p by RIP3 in the necrosome |
| **P17** | Association of CYLD to RIP1-Ub in complex I | **P37** | Phosphorylation of RIP3 by RIP1-p in the necrosome |
| **P18** | Dissociation of CYLD from complex I | **P38** | Association of MLKL to the necrosome |
| **P19** | Deubiquitination of RIP1-Ub by A20 in complex I | **P39** | Dissociation of MLKL from the necrosome |
| **P20** | Deubiquitination of RIP1-Ub by CYLD in complex I | **P40** | Phosphorylation of MLKL |

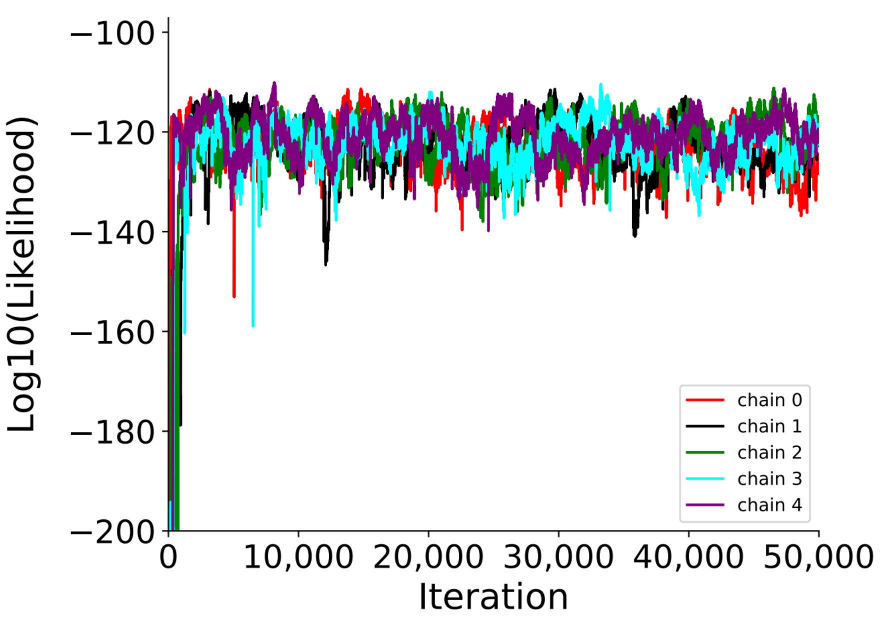


**Supplementary Figure S1. Log-likelihood vs. iteration for all five Markov chains used in the Bayesian parameter calibration.** For each chain, the first 25,000 iterations were discarded (considered burn-in), leaving a total of 125,000 parameter sets total, of which 10,628 are unique.

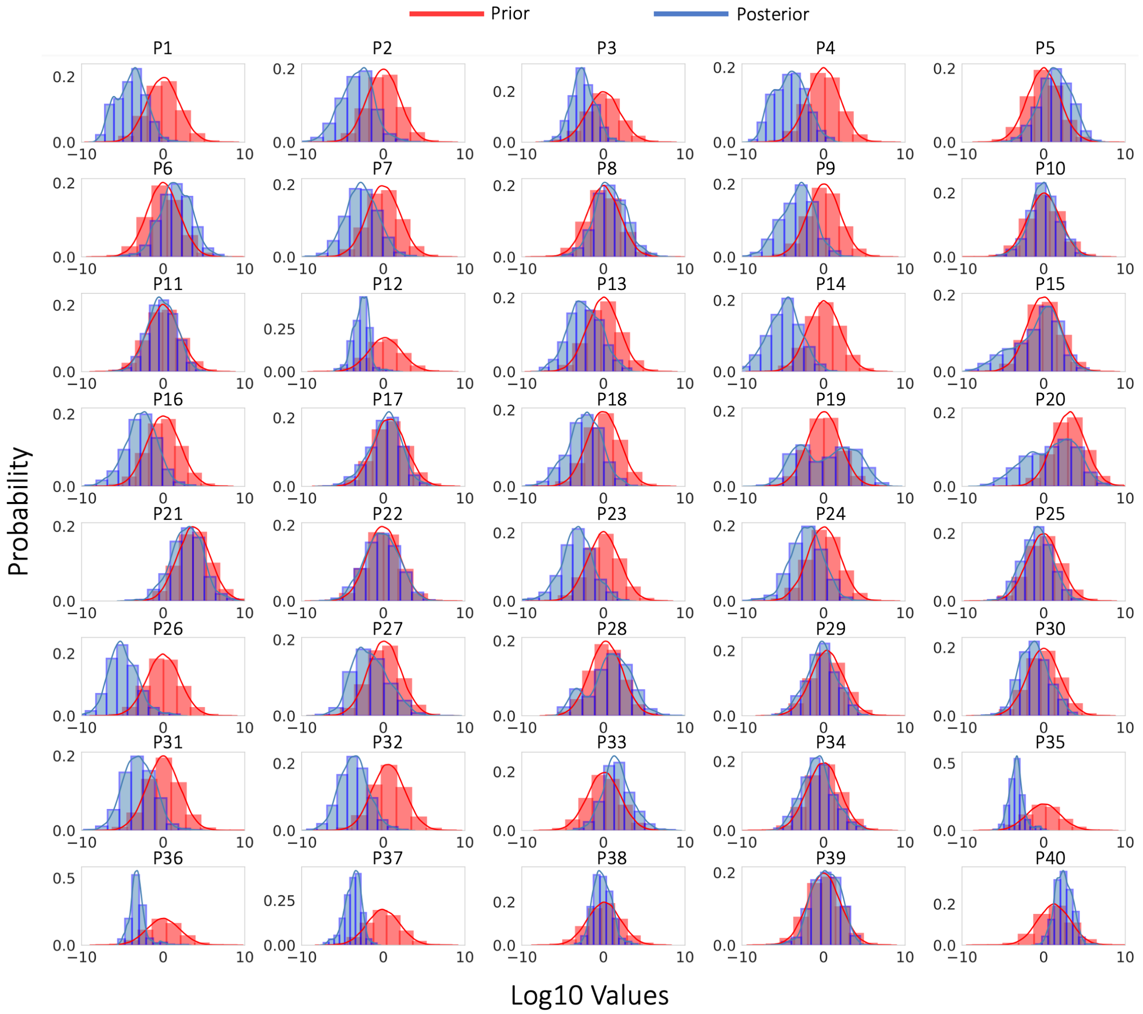


**Supplementary Figure S2. Distributions of parameter values from Bayesian model calibration.** Both prior (red) and posterior (blue) distributions are shown.

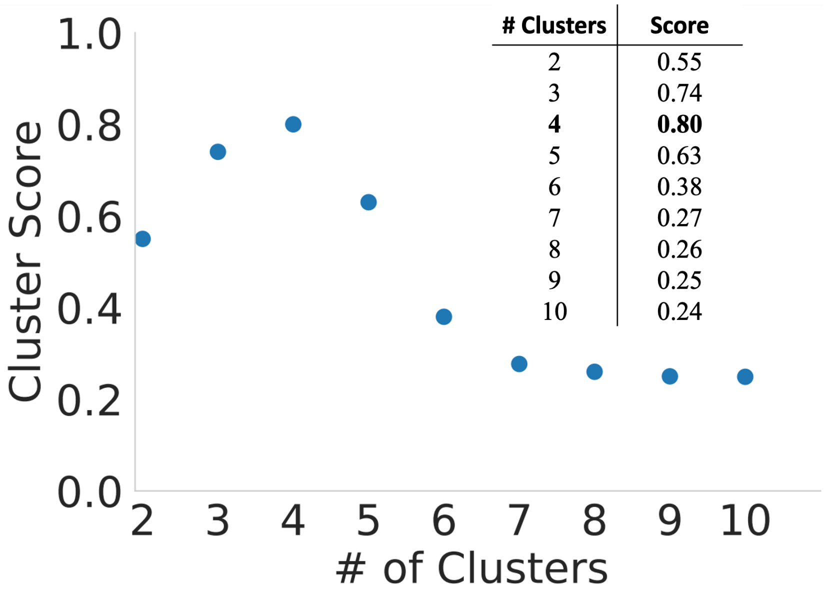


| # Clusters | Score |
|---|---|
| 2 | 0.55 |
| 3 | 0.74 |
| **4** | **0.80** |
| 5 | 0.63 |
| 6 | 0.38 |
| 7 | 0.27 |
| 8 | 0.26 |
| 9 | 0.25 |
| 10 | 0.24 |

**Supplementary Figure S3: Silhouette clustering scores for determining the number of modes of necroptosis execution.** The maximum value is for four clusters. Values were also calculated for 11-20 clusters and were all <0.3 (data not shown).

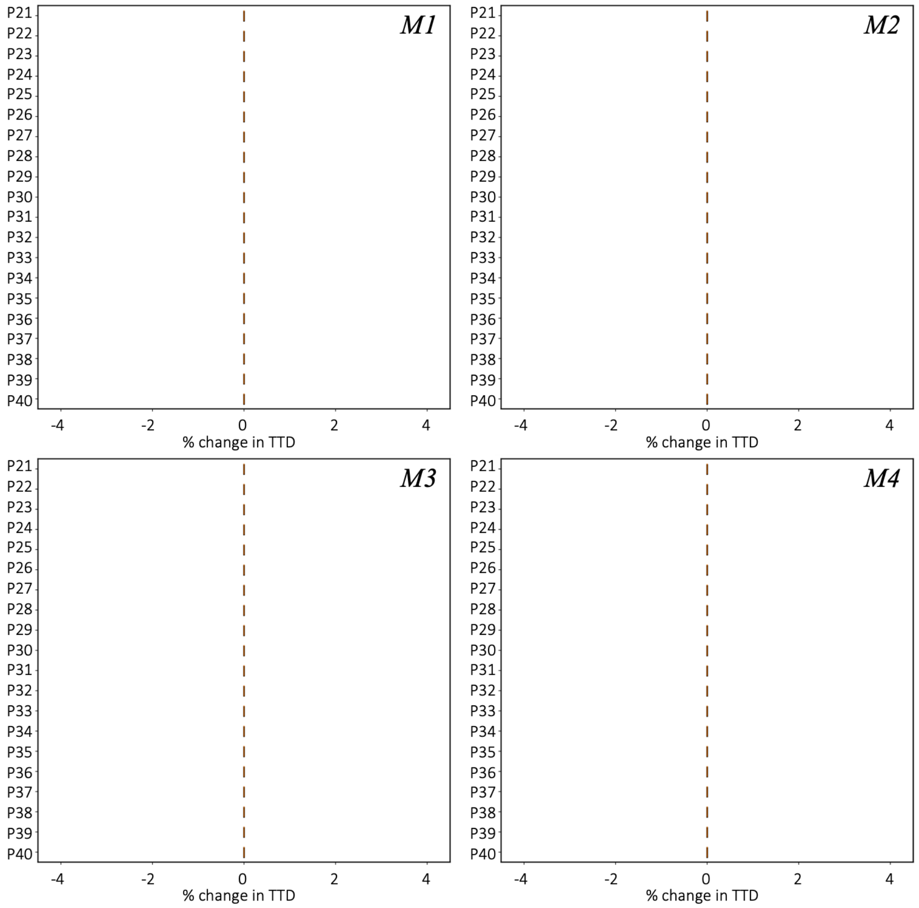


**Supplementary Figure S4: Rate constant sensitivity analyses show no sensitivity for rate constants P21-P40 in any mode.** Values were varied in a range ± 20% around the reference parameter set for each mode.

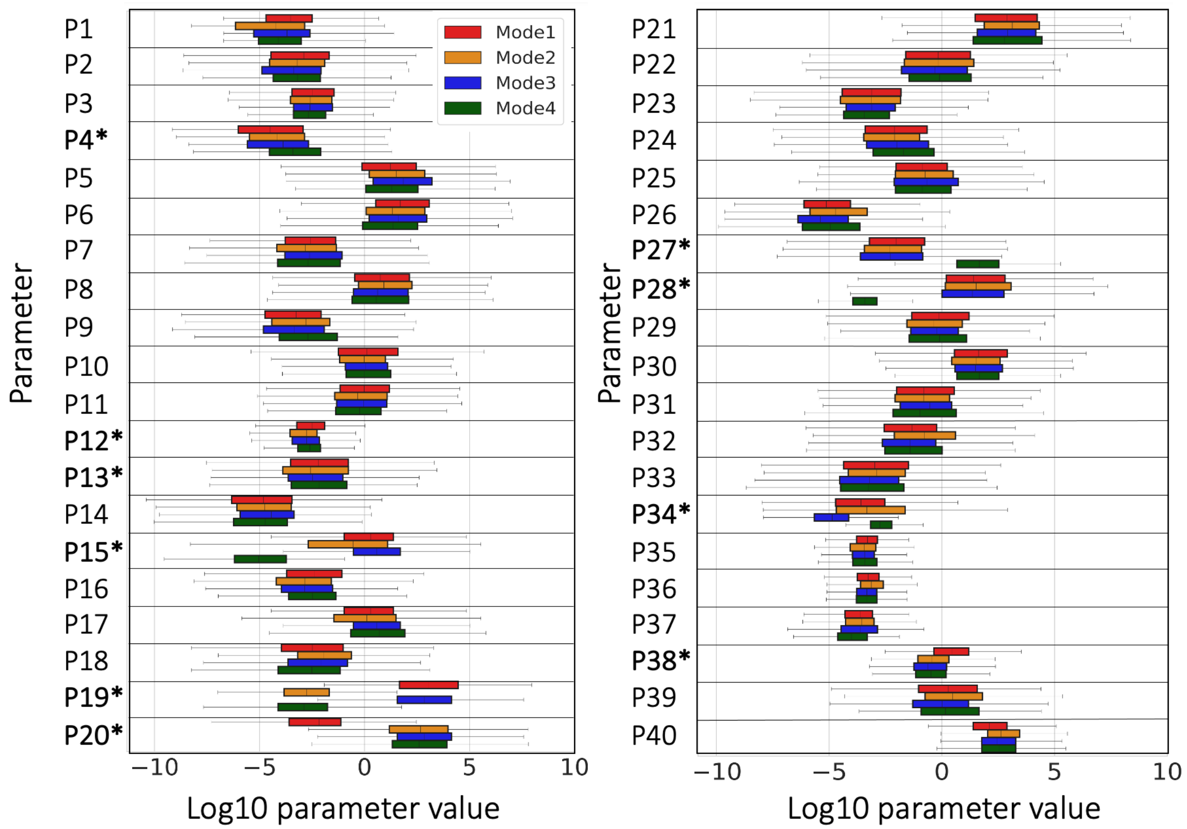


**Supplementary Figure S5: Rate constant distributions for all four modes of execution.** Parameters denoted with an asterisks (*) are included in Fig. 2C of the main text.

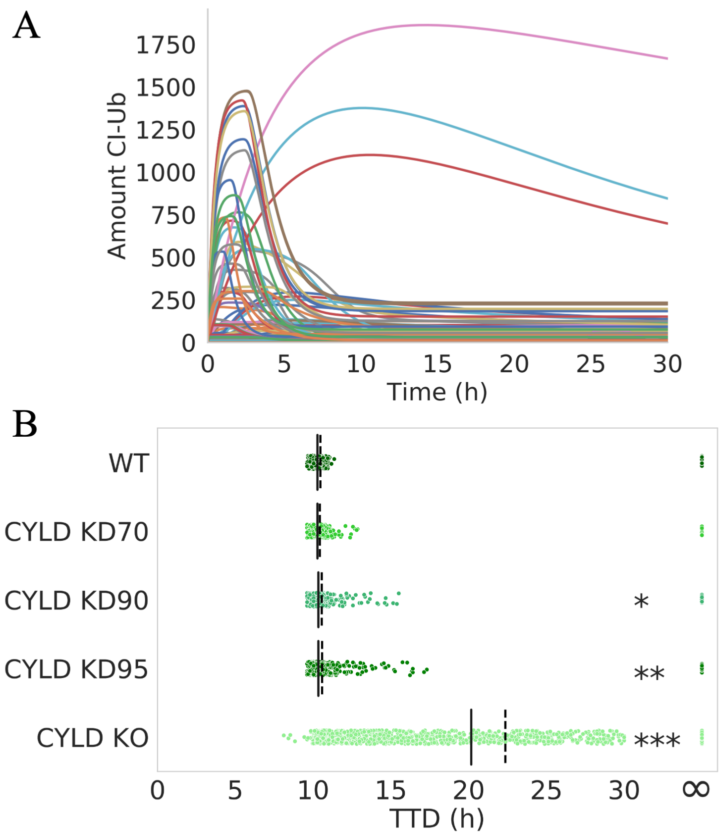


**Supplementary Figure S6: Dynamics in necroptosis execution mode 4.** (A) Time courses for ubiquitinated complex I for all parameter sets in mode 4 show that CYLD (9,075 molecules; Supplementary Table S1) is always in great excess. (B) TTD distributions over all parameter sets in mode 4 for CYLD knockdowns (KDs; 70%-95%) and knock out (KO), compared to wild-type (WT). Solid black lines = medians, dashed black lines = means; * $p < 0.05$, ** $p < 0.01$, *** $p < 0.001$ (Mood's median test).
